# Optimal implementation of genomic selection in clone breeding programs - exemplified in potato: I. Effect of selection strategy, implementation stage, and selection intensity on short-term genetic gain

**DOI:** 10.1101/2022.11.25.517496

**Authors:** Po-Ya Wu, Benjamin Stich, Juliane Renner, Katja Muders, Vanessa Prigge, Delphine van Inghelandt

**Affiliations:** Institute of Quantitative Genetics and Genomics of Plants, Heinrich Heine University, 40225 Düsseldorf, Germany; Cluster of Excellence on Plant Sciences (CEPLAS), Heinrich Heine University, 40225 Düsseldorf, Germany; Max Planck Institute for Plant Breeding Research, 50829 Köln, Germany; Böhm-Nordkartoffel Agrarproduktion GmbH & Co. OHG, 17111 Hohenmocker, Germany; NORIKA GmbH, 18190 Sanitz, Germany; SaKa Pflanzenzucht GmbH & Co. KG, 24340 Windeby, Germany

**Author notes:** Corresponding author: Delphine van Inghelandt.

**Keywords:** Clone breeding, Genetic gain, Genomic selection, Optimum allocation, Potato, Selection strategy

## Abstract

Genomic selection (GS) is used in many animal and plant breeding programs to enhance genetic gain for complex traits. However, its optimal integration in clone breeding programs that up to now relied on phenotypic selection (PS) requires further research. The objectives of this study were to (i) investigate under a fixed budget how the weight of GS relative to PS, the stage of implementing GS, the correlation between an auxiliary trait assessed in early generations and the target trait, the variance components, and the prediction accuracy affect the genetic gain of the target trait of GS compared to PS, (ii) determine the optimal allocation of resources maximizing the genetic gain of the target trait in each selection strategy and for varying cost scenarios, and (iii) make recommendations to breeders how to implement GS in clone and especially potato breeding programs. In our simulation results, any selection strategy involving GS had a higher short-term genetic gain for the target trait than Standard-PS. In addition, we show that implementing GS in consecutive selection stages can largely enhance short-term genetic gain and recommend the breeders to implement GS at single hills and A clone stages. Furthermore, we observed for selection strategies involving GS that the optimal allocation of resources maximizing the genetic gain of the target trait differed considerably from those typically used in potato breeding programs. Therefore, our study provides new insight for breeders regarding how to optimally implement GS in a commercial potato breeding program to improve the short-term genetic gain for their target trait.

## INTRODUCTION

Potato (*Solanum tuberosum* L.) is with respect to the production volume one of the most important food crops in the world after sugarcane, maize, wheat, and rice (FAO-STAT (2020), http://www.fao.org/faostat/en/). However, in contrast to other crops, only a low genetic gain was observed for yield in the past decades (Stokstad, 2019; Ortiz et al., 2022). The selection gain is, compared to the one in homozygous diploid species, limited by the high heterozygosity and tetraploidy of potato. (Lindhout et al., 2011; Jansky et al., 2016). In addition, potato has a low multiplication coefficient (Grüneberg et al., 2009), which leads to the availability of only one or few tubers per genotype for phenotypic evaluation at early stages in the breeding program (Gopal, 2006). This delays the evaluation of traits related to productivity (such as tuber yield) or quality, as they rely on multi-location field trials and/or destructive assessment, and these can only be performed after one to several multiplication steps. As a consequence, only traits which can be assessed based on a low number of plants can be considered in the early stages of potato breeding programs. In contrast, target traits whose evaluation requires many plants and/or environments can only be selected for in later stages of the breeding program. However, the correlation between the early measured and target traits is variable and can be very low. Furthermore, the evaluation of target traits in potato is more expensive compared to their evaluation in non-clonal crops as a considerably lower level of mechanization is currently possible. Therefore, clone and especially potato breeding programs would highly benefit from the possibility to select for target traits at early stages of the breeding program e.g. with the implementation of genomic selection (GS).

GS proved to enhance genetic gain for complex traits in both animal and plant breeding programs (Meuwissen et al., 2001; Desta and Ortiz, 2014). This is because GS allows to predict the performance of target traits without phenotypic evaluation in early stages. The selection on target traits at early stages using estimated genetic values (EGV) avoids discarding those individuals with desirable alleles for the trait, which will increase the genetic gain per year. In addition, the performance prediction of target traits without phenotypic evaluation in early stages has the potential to reduce the length of the breeding cycle. One parameter that influences the potential of GS is the prediction accuracy.

Several empirical studies have explored the potential of implementing GS in potato breeding for different traits by determining the prediction accuracy (Slater et al., 2016; Sverrisdóttir et al., 2017; Enciso-Rodriguez et al., 2018; Endelman et al., 2018; Stich and Van Inghelandt, 2018; Sverrisdóttir et al., 2018; Caruana et al., 2019; Byrne et al., 2020; Gemenet et al., 2020; Sood et al., 2020; Wilson et al., 2021). Different degrees of prediction accuracies from low to high depending on the studied traits have been reported, which could be caused by the different genetic architectures, prediction models, but also the considered genetic material. However, only few studies evaluated the effect of GS on the genetic gain for the studied traits. One of them was Slater et al. (2016), who estimated that the genetic gain after implementing GS for complex traits was higher than that of PS. The results of Stich and Van Inghelandt (2018) suggested that for some traits GS leads to a higher gain of selection than PS even without reducing the cycle length. However, no earlier study considered directly the aspect that PS and GS need to be compared at a fixed budget. Furthermore, when implementing GS in a clone breeding program, the selected proportion of PS on the early trait will be partially shifted to GS on the target trait. This shift can be realized to different degrees and the resulting selected proportion for PS or GS might influence the efficiency of the selection strategy. Therefore, for the implementation of GS in clone breeding programs not only the prediction accuracy of the GS model but also its relative weight to PS has to be examined. Furthermore, these aspects are influenced by the correlation between the early and the target trait and also the variance components of the considered trait have an influence on the genetic gain. However, the influences of these parameters and their interaction on the genetic gain in clone breeding programs have not been investigated until now.

Werner et al. (2020) investigated different strategies to implement GS in clone breeding programs exemplarily with genome parameters of strawberry. They evaluated the performance of a breeding program that introduced GS in the first clonal stage and mainly focused on how to select parents for the next crosses and drive population improvement to enhance long-term genetic gain. However, in a classical clone breeding program, there are several stages where GS could be implemented and their effect on the gain of selection have not been studied so far.

Another aspect that needs to be decided during the implementation of GS in clone or potato breeding programs is the number of stages in which GS is applied. Once the clones are genotyped for the first GS application, the possibility of re-using the same EGV to perform GS in two or more stages is given. A similar idea was proposed by Spindel et al. (2015) for a rice breeding program but has neither been assessed by theoretical considerations nor by computer simulations nor any empirical experiments. To the best of our knowledge, no earlier study has investigated at which stage and in how many selection stages GS should be implemented in clonal crops to maximize the short-term genetic gain under a given budget.

Optimum allocation of resources under a given budget is essential to improve the efficiency of breeding programs (Longin et al., 2006). However, most studies on the implementation of GS in breeding programs neglected this effect. Longin et al. (2015) and Marulanda et al. (2016) assessed this point for cereal breeding programs. However, to the best of our knowledge, no earlier study is available about the effect of the implementation of GS on the optimum allocation of resources in clone breeding programs.

The objectives of this study were to (i) investigate under a fixed budget how the weight of GS relative to PS, the stage of implementation of GS, the correlation between traits (auxiliary trait assessed in early generations and target trait), the variance components, and the prediction accuracy affect the short-term genetic gain of the target trait in potato breeding programs compared to PS, (ii) determine the optimal allocation of resources maximizing the short-term genetic gain of the target trait in each selection strategy and for varying cost scenarios, and (iii) make recommendations to breeders how to implement GS in clone breeding programs.

## MATERIALS AND METHODS

### Empirical basis of the computer simulations

Our simulations were based on an empirical genomic dataset of tetraploid potato. This empirical genomic dataset comprised 19,649,193 sequence variants revealed in a diversity panel of 100 tetraploid potato clones (Baig et al. in preparation). The unphased sequence variants included single nucleotide polymorphism (SNP) and insertion/deletion (InDel) polymorphisms. Sequence variants with a minor allele frequency < 0.05 and missing rate > 0.1 were removed. The 100 clones were used as parents of the simulated progenies and will be called parental clones hereafter.

The progenies were simulated using AlphaSimR (Gaynor et al., 2021). For this, the genetic map information of all genomic variants was estimated using a Marey map (for details see Method S1 and Figure S1). Subsequently, the genomic information for each variant served as input for the simulations.

### Simulation of initial population

To stick to the size of commercial breeding programs (Breeders personal communication, Table 1) an initial population of 300,000 clones was simulated like described here under. From all possible crosses in the half-diallel among the 100 parental clones, 300 were randomly selected. For each of these 300 crosses, 1,000 F1 progenies were simulated using AlphaSimR. The two steps of this procedure (the random selection of 300 crosses and the simulation of their progenies) was repeated 1,000 times independently.

**Table 1:** Dimensioning of a standard potato breeding program that exclusively relies on phenotypic selection.

| Stage | Number of clones | Number of locations | Phenotyping cost per clone and plot (€) | Cost per stage (€) |
| --- | --- | --- | --- | --- |
| Seedling | 300,000 | 1 | 1.4 | 420,000 |
| Single hills | 100,000 | 1 | 1.4 | 140,000 |
| A clone | 10,000 | 1 | 1.4 | 14,000 |
| B clone | 1,500 | 2 | 25 | 75,000 |
| C clone | 300 | 3 | 25 | 22,500 |
| D clone | 60 | 4 | 25 | 6,000 |
| Sum |  |  |  | 677,500 |

### Simulation of true genetic and phenotypic values Target trait (T_t_)

In our study, a genetically complex target trait representing the weighted sum of all market relevant quantitative traits was considered and will be named T_t_ hereafter. A random set of 2,000 sequence variants were considered as quantitative trait loci (QTL) for T_t_. The true additive effects of the 2,000 QTL were drawn from a gamma distribution (cf. Hayes and Goddard, 2001) with *k* = 2 and *θ* = 0.2, where *k* and *θ* are shape and scale parameter, respectively. To control the degree of dominance *δ* between 0 and 1 for each QTL, the ratios of dominance to additive effect were produced from a beta distribution with the two shape parameters *a* = 2 and *b* = 2. The true dominance effect at each QTL was then calculated by multiplying the true additive effect by the QTL specific *δ* (Figure S2). For each QTL, all possible genotype classes were AAAA, AAAB, AABB, and BBBB, which were respectively coded from 0, 1, 2, 3, and 4 for additive effect; and 0, 1, 1, 1, and 0 for dominance effect. Finally, the true genetic value for T_t_ (TGV_T_t__) was calculated for each clone by summing up the true additive and dominance effects at the 2,000 QTL.

In order to simulate phenotypic values, two ratios of variance components (VC) were assumed for 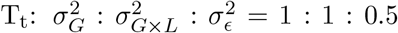 (VC1) and 1: 0.5: 0.5 (VC2), where 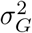 denoted the genotypic variance, 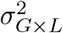 the variance of interaction between genotype and location, and 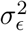 the error variance. The genotypic variance was estimated by the sample variance of TGV_T_t__ in the initial population. The phenotypic value for the target trait was then calculated as P_T_t__ = TGV_T_t__ + *ϵ*_T_t__, where *ϵ*_T_t__ was the non-genetic value following a normal distribution 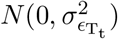, with

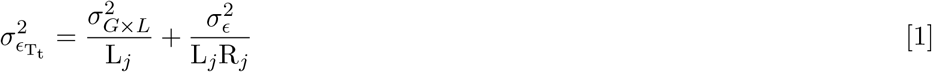

representing the non-genetic variance, in which *L_j_* was the number of locations at stage *j*, and *R_j_* the number of repetitions at stage *j*. We set the number of replications to one (*R_j_* = 1) in each location (cf. Melchinger et al., 2005).

### Phenotypic trait assessed in early generations of the breeding program (*T_a_*)

The weighted sum of the auxiliary traits measured in the first three generations of the breeding program will be referred to as T_a_ hereafter. To control the genetic correlations between T_a_ and T_t_ (r), the true genetic values for T_a_ were generated by TGV_T_a__ = TGV_T_t__ + *ϵ*_r_, where *ϵ*_r_ was the residual value following a normal distribution 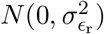, with

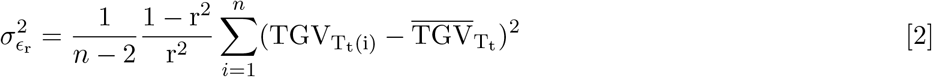

determined by the degree of r, where *n* was the number of clones for the initial population, TGV_T_t_(i)_ the TGV for T_t_ of the *i^th^* clone, and 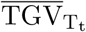 the average of TGV_T_t__ in the initial population. Then, the phenotypic value for T_a_ was calculated as P_T_a__ = TGV_T_a__ + *ϵ*_T_a__, where *ϵ*_T_a__ was a non-genetic value following a normal distribution 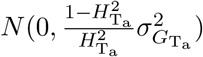, in which 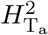 was the broad-sense heritability for T_a_, and 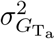 the genetic variance of T_a_ and estimated by the sample variance of TGV_T_a__ in the initial population. In this study, 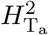 was set as 0.6.

### Simulation of estimated genetic values

In this study, we assumed that a GS model for T_t_ with a prediction accuracy of PA was available. The estimated genetic values of T_t_ obtained from the GS model were estimated by EGV_T_t__ = TGV_T_t__ + *ϵ*_PA_, where *ϵ*_PA_ was the residual value following a normal distribution 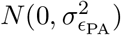, with

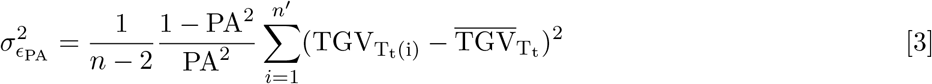

determined by the level of PA, where *n*′ was the number of genotyped clones (= N_GS_), TGV_T_t_(i)_ the TGV of the target trait at the *i^th^* genotyped clone, and 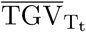 the average of TGV_T_t__ on all N_GS_ genotyped clones.

### Selection strategies

#### Standard breeding program

A standard potato breeding program relying exclusively on PS (Standard-PS) was considered as benchmark (Figure 1). To simplify the comparison between PS and GS strategies, we considered in this study six testing stages in the potato breeding program. The six testing stages were seedling, single hills, and A, B, C, and D clone stages, abbreviated in the following as SL, SH, A, B, C, and D, respectively. The number of tested clones (N) and locations (L) for each testing stage are shown in Table 1. The selected proportions from SL to SH (p_1_), SH to A (p_2_), A to B (p_3_), B to C (p_4_), and C to D (p_5_) were set to 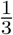, 0.1, 0.15, 0.2, and 0.2, respectively, as estimates from typical commercial potato breeding programs (Breeders personal communication). The selection in the early stages (SL, SH, and A) was based on the phenotypic value of the auxiliary trait P_T_a__, and for the late stages (B, C, and D) on the phenotypic value of the target trait P_T_t__(Figure 1).

**Figure 1:**
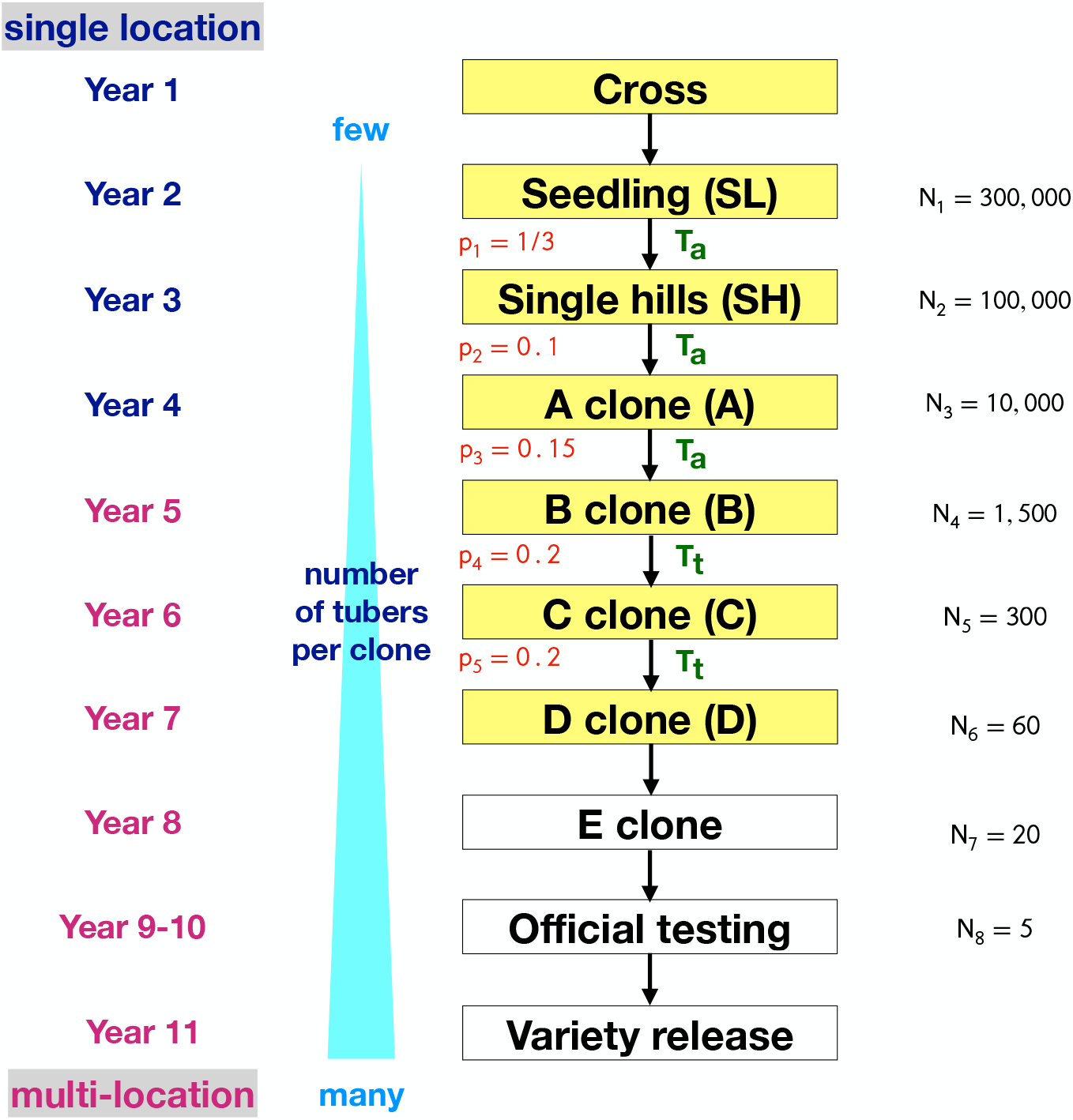
The standard clone breeding program examined in this study that relies exclusively on phenotypic selection. p_1_-p_5_ are the selected proportions from SL to SH, SH to A, A to B, B to C, and C to D, respectively, where SL, SH, A, B, C, and D represent the stages of seedling, single hills, A, B, C, and D clones. T_a_ represented the integral of early measured traits and T_t_ the integral of the target traits. The yellow marked stages are those that were examined in our study.

#### Breeding programs involving genomic selection

Three GS strategies were evaluated in which GS was implemented at the (1) seedling, (2) single hills, and (3) A clone stage, abbreviated as GS-SL, GS-SH, and GS-A, respectively. All selection steps of the GS strategies were similar to those of the standard breeding program except the following modifications (Figure 2). Here, the strategy GS-SL will be taken as an example for the description. In the seedling stage, N_1_ clones were evaluated for P_Ta_. From these N_1_ clones, the N_GS_ ones with a higher P_Ta_ were genotyped. *α*_1_ was defined as ratio of N_GS_ to N_1_, i.e. the proportion of clones selected by PS to be genotyped. Then, N_2_ clones were selected based on the EGV_Tt_ in the N_GS_ genotyped clones for the single hills stage. Afterwards, the selection process in the following stages were the same as Standard-PS. For the other two GS strategies, GS-SH and GS-A, the selection was performed accordingly. For each stage *k* in which GS was applied, the corresponding *α_k_* was larger than p_k_, where 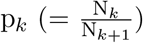 was the selected proportion between the two stages to which GS was applied. *k* was set to 1, 2, and 3 for the strategies (1) GS-SL, (2) GS-SH, and (3) GS-A, respectively (Figure 2).

**Figure 2:**
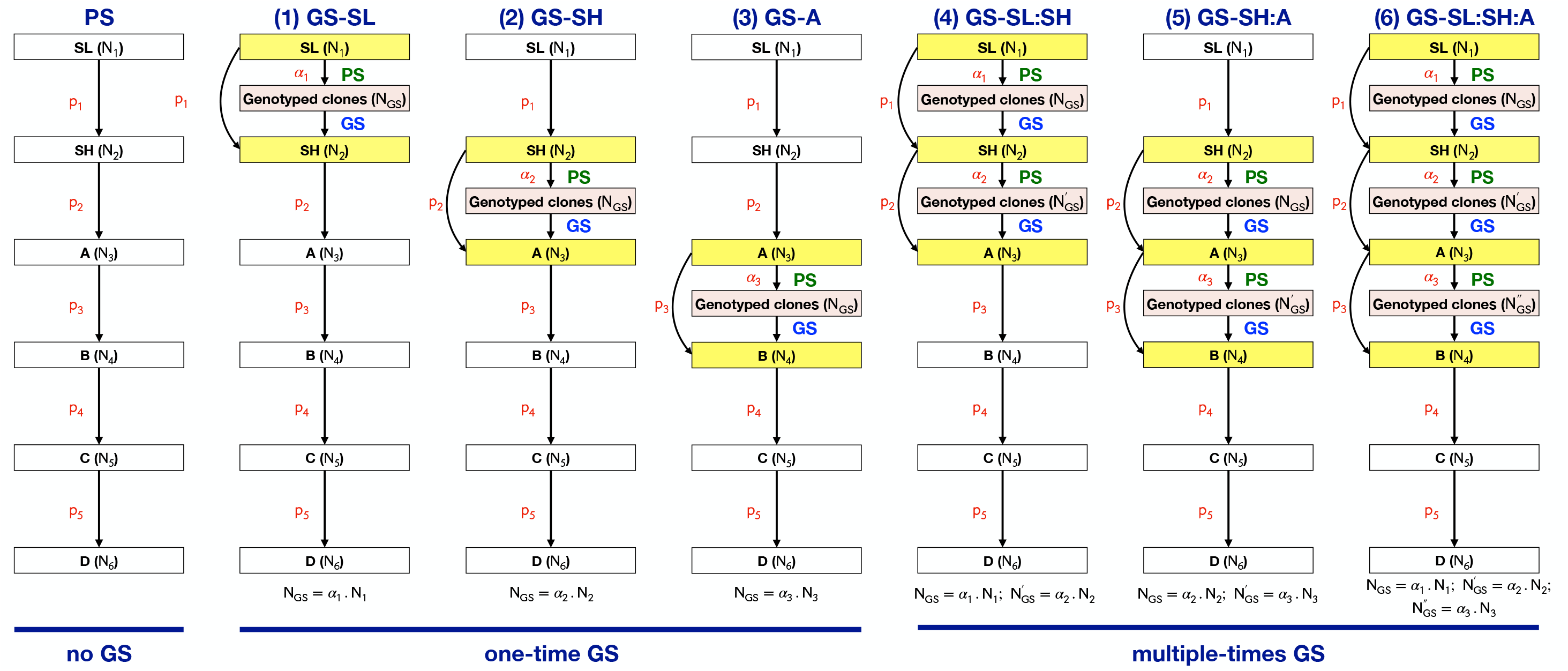
Graphical illustration of the standard as well as the six selection strategies that include genomic selection that were examined in our study. p_1_-p_5_ are the selected proportions from SL to SH, SH to A, A to B, B to C, and C to D, respectively, where SL, SH, A, B, C, and D represent the stages of seedling, single hills, A, B, C, and D clones. *α_k_* the proportion of clones selected by PS to be genotyped in stage *k* and N_*k*_ is the number of clones of the respective stage.

To evaluate whether adopting the same GS model for selection on T_t_ in several stages improves the short-term genetic gain compared to using GS only once, we evaluated three additional strategies (Figure 2):

(4) GS-SL:SH – GS was applied not only at seedling stage but also at single hills stage;

(5) GS-SH:A – GS was applied not only at single hills stage but also at A clone stage; and

(6) GS-SL:SH:A – GS was applied at seedling, single hills and A clone stages.

For these three GS strategies, genotyping of NGS clones only took place when GS was used for the first time. When GS was used a second or third time, the same EGV_T_t__ for the tested clones from the initial GS model were used for the selection.

### Economic settings and additional quantitative genetic parameters

In this study, the costs for phenotypic evaluation of T_a_ and T_t_ in one environment were assumed to be 1.4 and 25 €, respectively. The costs for genotypic evaluation per clone were assumed as 25 €(Table 1). To compare the short-term genetic gain of T_t_ (Δ*G*) between Standard-PS and several GS strategies, the budget across different selection strategies was fixed to 677,500 €. Therefore, the number of tested clones in seedling stage (N_1_) must be adjusted/reduced when introducing GS into a breeding program to compensate for the additional genotyping cost. In the first part of the simulations, the selected proportions were fixed to those of Standard-PS. This was realized in our study by randomly sampling the reduced N1 from the initial population with an equal sample size for each cross population.

We were interested in how different values of r, PA, VC, and L influence Δ*G*. Therefore, three different levels of r (−0.15, 0.15 and 0.3), PA (0.3, 0.5 and 0.7), and two different ratios of VC for T_t_ (see above) were examined in our simulations. The selection of clones based on T_t_ that was assessed in field experiments in more than one location happened at B and C clone stages. Thus, we varied the number of locations from 2 to 4 and 3 to 6 in increments of 1 for B and C clone stages, respectively, and designated them as L_4_ and L_5_. Furthermore, to investigate how different levels of *α_k_* affect Δ*G*, we varied *α_k_* from 0.4 to 0.9 in increments of 0.1 for the strategies GS-SL, GS-SL:SH, and GS-SL:SH:A, and from 0.2 to 0.9 in increments of 0.1 for the other strategies. Δ*G* was calculated as the difference in mean genetic values of T_t_ between the D clone and the seedling stage (cf. Longin et al., 2015; Marulanda et al., 2016).

### Optimum allocation of resources

In the below described simulations, we relaxed the restrictions of the above described simulations that the selected proportions were fixed to those of Standard-PS. To determine the optimum allocation of resources maximizing Δ*G* under a given budget, a general linear cost function to aggregate all costs across all stages in the breeding program was created:

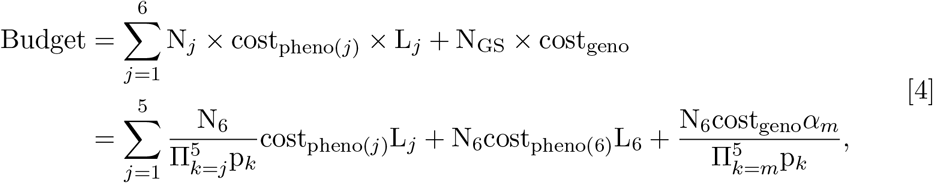

where *N_j_* was the number of clones at stage *j*, cost_pheno(*j*)_ the cost for phenotypic evaluation at stage *j*, N_GS_ the number of genotyped clones, and cost_geno_ the genotyping cost (for details see Method S2). In addition, p_k_ was the selected proportion from stage *j*(*m*) to stage *j*(*m*) + 1, where *m* was the stage in which GS was applied first. For more details, *m* = 1 referred to GS-SL, GS-SL:SH and GS-SL:SH:A; *m* = 2 for GS-SH and GS-SH:A; and *m* = 3 for GS-A. The GS strategies with optimum allocation of resources will be named Optimal-GS hereafter.

The optimum allocation was determined by a grid search across the permissible space of p_2_ to p_5_ and *α_k_* for a set of given input parameters. The latter included the number of tested clones at D clone stage (N_6_), the GS strategy, the phenotyping and genotyping costs, L, r, VC of T_t_, 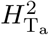, and the total budget. We set N_6_ to 60. In the grid search, any p_k_ varied between 0.1 and 0.5 in increments of 0.05 to avoid too strong/weak selections. *α_k_* was chosen as described above. Consequently, in each permissible allocation, p_1_ was completely determined by equation [4] under the constrained budget and the given input parameters. Subsequently, the mean genetic gain across 1,000 simulation runs was calculated for each permissible allocation of the grid search. To obtain reliable estimates of the optimal allocation of resources, we performed a least significant difference (LSD) test on Δ*G* across all permissible allocations of the grid search within a specific scenario. We selected the significant group showing the maximum Δ*G* among all permissible sets and then considered the average of the allocations as optimal result.

The above described simulations required for some grid search sets (those with low p_1_ to p_3_ but high p_4_ and p_5_) with more than 300,000 clones in the seedling stage. Thus, the size of the initial population was increased to 900,000 clones.

To investigate whether an increase of phenotyping cost of T_a_ and the genotyping cost have an influence on the optimal allocation of resources, we considered three different phenotyping costs for T_a_ (0.7, 1.05, and 1.4 €), and three different genotyping costs (15, 25, and 40 €).

## RESULTS

The mean genetic gain (Δ*G*) and genetic variance 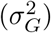 of the target trait at D clone were assessed considering different values of r, PA, *α_k_*, as well as different selection strategies. To easily compare among the examined strategies, the budget, the selection proportion between stages p_1_-p_5_ and the number of test locations were fixed according to those of the Standard-PS strategy.

Increasing r and PA either individually or simultaneously led to a higher Δ*G* (Figure 3 and S4). Regardless of PA and r, any selection strategy incorporating GS was superior to the Standard-PS strategy with respect to Δ*G* (Figure 3). Low or negative values for r and high PA increased this tendency even more. The least improvement of Δ*G* relative to Standard-PS was observed across all scenarios for the strategy GS-SL. The strategies GS-A and GS-SH resulted in considerably higher values for Δ*G* relative to PS and under the scenarios with low r but high PA, the latter strategy was significantly superior to the former.

**Figure 3:**
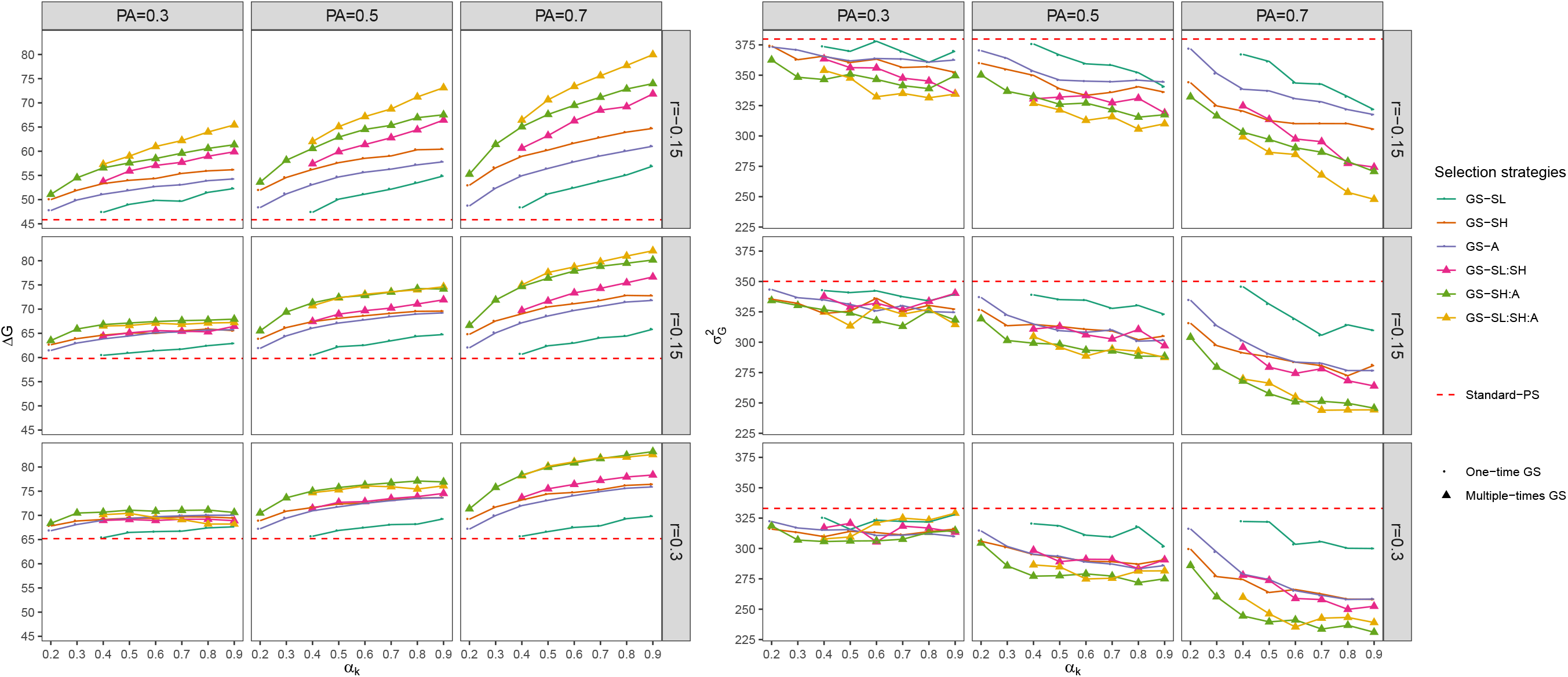
Genetic gain (Δ*G*, left) and genetic variance (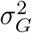, right) for the target trait on average across 1,000 simulation runs at D clone stage for different weights of genomic selection (GS) relative to phenotypic selection (*α_k_*), different selection strategies, different correlations between the traits (r=-0.15, 0.15, and 0.3), prediction accuracies (PA=0.3, 0.5, and 0.7), and for the ratio of variance components VC1 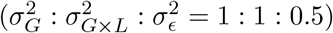. The details regarding the selection strategies are shown in Figure 2.

Implementing GS in successive stages (i.e. GS-SL:SH, GS-SH:A, and GS-SL:SH:A) had an advantage over the strategies using GS one time, except for the scenario with the lowest PA (=0.3) but the highest r (=0.3). The ranking of performance among these strategies was GS-SL:SH:A > GS-SH:A > GS-SL:SH. The difference among these strategies was lower, if r increased or PA decreased.

For all GS strategies, higher *α_k_* values led to reductions in the number of clones available in the seedling stage (Figure S3), but increased Δ*G* (Figure 3). For all except eight scenarios, the highest Δ*G* was observed if *α_k_* was at its maximum (0.9). The remaining scenarios in which the maximum Δ*G* were observed for *α_k_* =0.7 or 0.8 instead of 0.9, however, showed Δ*G* values that were not significantly different from the Δ*G* values observed for *α_k_* =0.9 (data not shown). Only for GS-SL:SH:A an exception was observed from this trend, namely that the maximal Δ*G* was observed for *α_k_*=0.5 for the scenario with r=0.3 and PA=0.3. In accordance to the above described observations regarding the differences among selection strategies, also the differences among Δ*G* for the different levels of *α_k_* were low for the scenarios with high r and/or low PA.

In all the above described simulations of the selection strategies that exploit GS in several stages, *α_k_* was the same for each stage in which GS was applied. However, for these strategies, we also evaluated whether varying *α_k_* had an influence on Δ*G*. For the strategies GS-SL:SH and GS-SH:A, we observed that an increase of both *α_k_* values (i.e. *α*_1_ and *α*_2_ or *α*_2_ and *α*_3_) a higher Δ*G* was observed (Figure S5). The combination of two *α_k_* values that resulted in the highest Δ*G* was 0.84 and 0.79 or 0.86 and 0.86 for the respective strategies. A similar trend was observed for GS-SL:SH:A (Figure S6). However, for the scenarios with high r (=0.3), intermediate values of *α*_1_ were sufficient to result with high values of *α*_2_ and *α*_3_ in the maximal values of Δ*G* of 0.4-0.5 (Table S1).

The effect of variation of selection strategies, *α_k_*, r, and PA on the genetic variance were opposite to their effect on genetic gain (Figure 3). The scenarios with a higher genetic gain showed a lower genetic variance.

We also investigated the effects of different ratios of variance components (VC1 and VC2) and number of locations for phenotypic evaluation (L_4_ and L_5_) on Δ*G*. The ranking of the selection strategies with respect to Δ*G* was not affected by the studied ratios of VC (Figure 3 and S7). When 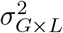 was halved (i.e. VC2 vs. VC1), Δ*G* increased from 3% to 8% depending on the selection strategies, PA, r, and *α_k_* (Figure S8). Although increasing L caused a decrease in the number of clones that are available at the seedling stage to compensate for additional phenotyping costs, Δ*G* significantly increased with increasing number of locations that were used for the evaluation of B and C clones (Figure 4). This trend was independent of selection strategies, PA, r, and *α_k_*. In all scenarios, the highest Δ*G* was observed with the highest number of locations in the B and C clone stages, i.e., L_4_ = 4 and L_5_ = 6. In these cases, Δ*G* was increased by 8% compared to Standard-PS with (L_4_, L_5_) = (2, 3).

**Figure 4:**
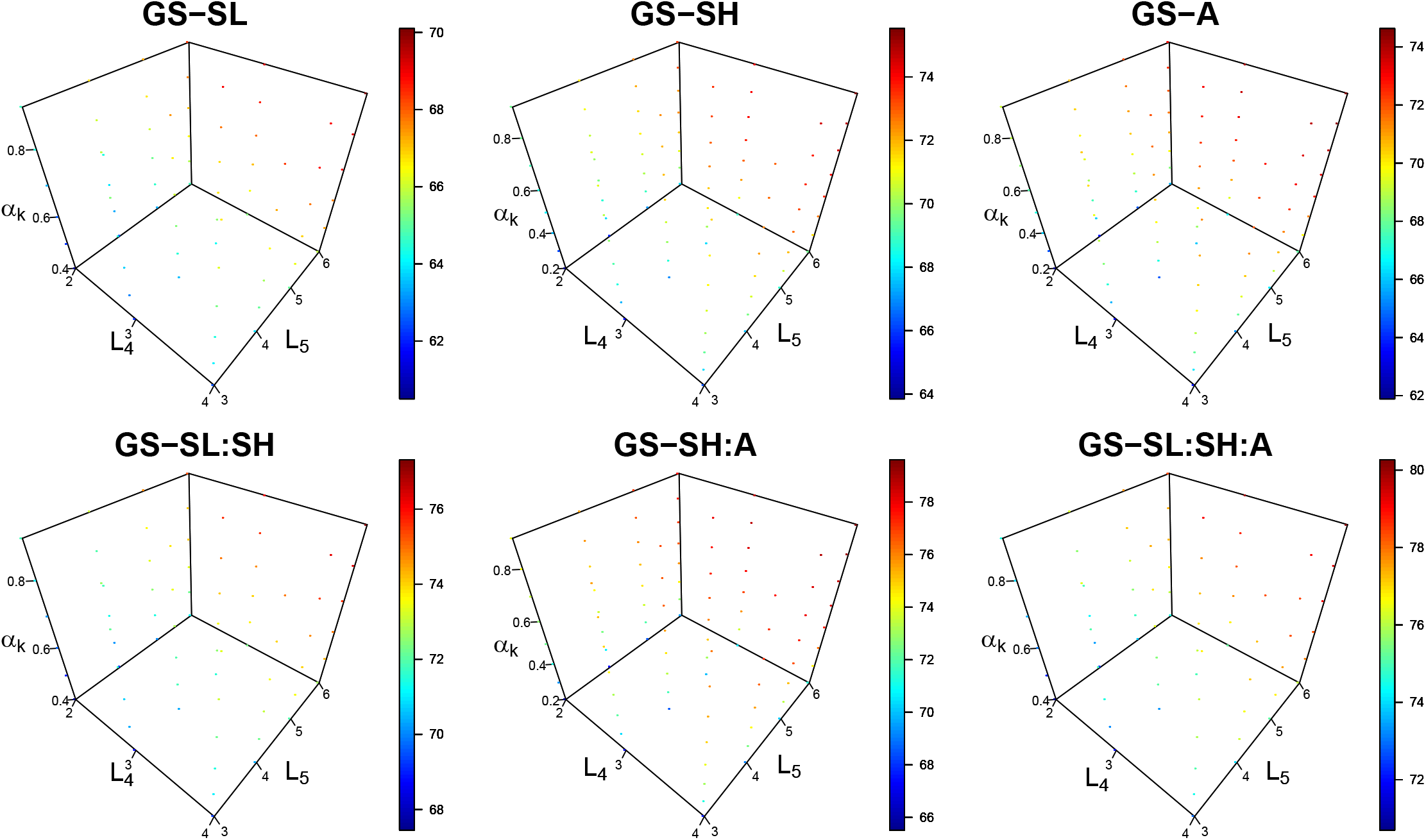
Genetic gain for the target trait (Δ*G*) on average across 1,000 simulation runs at the D clone stage for six different selection strategies with genomic selection (GS) for varying numbers of locations in the B and C clone stages (L_4_ and L_5_) and different weights of genomic selection (GS) relative to phenotypic selection (*α_k_*) when the correlation between the two traits was set to 0.15 and prediction accuracy was set to 0.5.

The optimal allocation of resources was assessed via a grid search across p_1_ - p_5_ and *α_k_*, *k* ∈ [1, 3] in a scenario with VC1, budget, L, and N_6_ like in the Standard-PS scenario. The optimum allocation of resources led also for the PS to an increase of Δ*G* (Optimal-PS) compared with the Standard-PS (Figure 5). On average across all evaluated scenarios, the strategy GS-SL had the worst performance out of the strategies incorporating GS. In a scenario with r < 0 and PA > 0.5, any selection strategy with GS revealed a higher Δ*G* than the Optimal-PS. The strategy GS-SL:SH:A only outperformed the other selection strategies if r=-0.15. In contrast, the strategy GS-SH:A or GS-A resulted in the highest Δ*G* if r was > −0.15. On average across all the examined scenarios, the strategy GS-SH:A resulted in the highest and most stable Δ*G* values.

**Figure 5:**
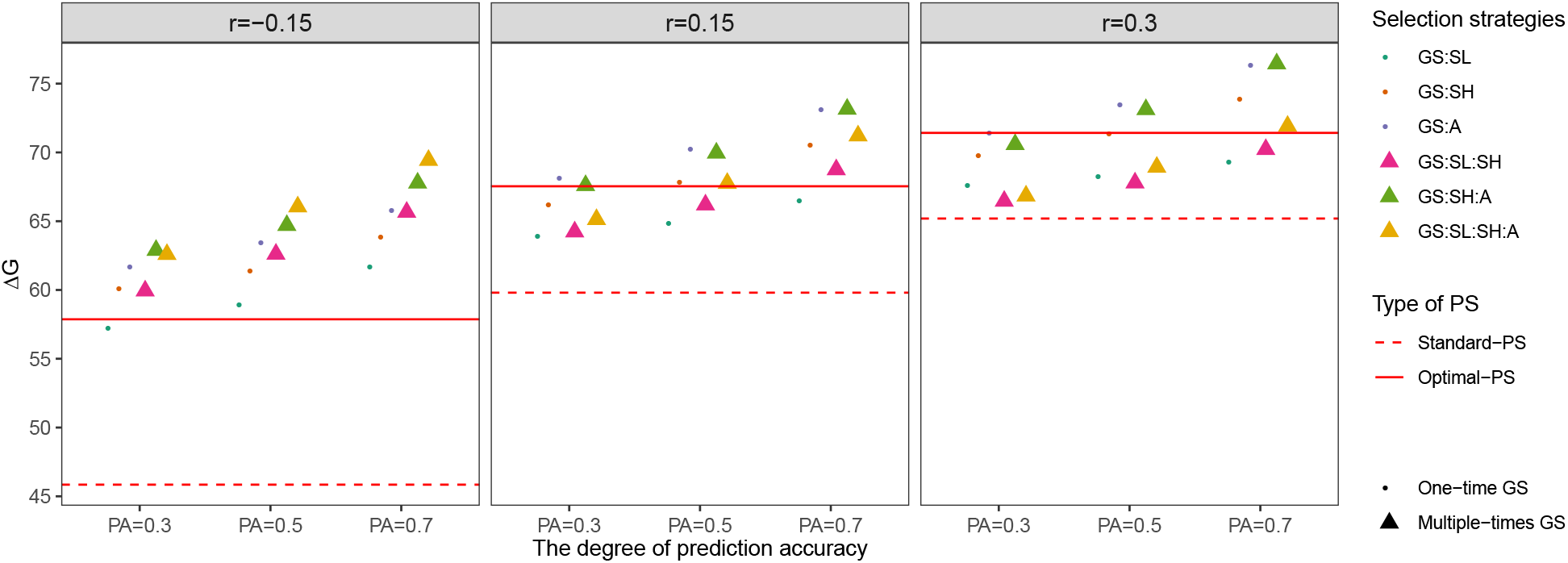
Genetic gain of the target trait (Δ*G*) after optimally allocated resources for different correlations between the traits (r=-0.15, 0.15, and 0.3) and different prediction accuracies (PA=0.3, 0.5, and 0.7). The presented Δ*G* values are the average of the genetic gains from the grid search sets that revealed no significant (*P* < 0.05) difference compared to the set with maximum genetic gain.

With the exception of one specific scenario, a high *α_k_* was required for each selection strategy to reach the maximal Δ*G* value (Table 2, S2 and S3). This exception was the strategy GS-SL in case of a positive r for which *α_k_* ranging from 0.21 to 0.61 resulted in the maximal Δ*G* values. Furthermore, to achieve maximum Δ*G* values, the selected proportions for the last two stages (i.e. p_4_ and p_5_) were low (0.17) on average across all scenarios. The level of the optimal p_*k*_ was influenced by the level of r as well as by the stage in which GS was implemented. In general, high optimal p_1_ values were observed with a negative correlation in comparison with the scenarios with a positive correlation. Furthermore, we observed for all strategies with implementation of GS that the selection proportion for that stage in which GS was applied was lower than the one observed at the same stage in the other strategies. This trend was more pronounced for scenarios with high PA. For instance, p_2_ (p_3_) for the strategy GS-SH (GS-A) was on average across all scenarios about 0.25 (0.21) lower than the one for the strategies excluding GS-SH (GS-A) with 0.42 (0.45).

**Table 2:** Optimum allocation of resources to maximize genetic gain of the target trait (Δ*G*) for the different selection strategies and correlations between the two traits (r=-0.15, 0.15, and 0.3). The prediction accuracy was 0.5 and the phenotyping cost of early measured trait 1.4 € and genotyping cost 25 €. p_1_ to p_5_, *α_k_*, and N_1_ are the selected proportion per stage, the weight of genomic selection relative to phenotypic selection, and the number of clones at the seedling stage, respectively. For description of selection strategies see text.

| Correlations | Selection strategies | $\Delta G^1$ | $SD_{\Delta G}^2$ | $p_1$ | $p_2$ | $p_3$ | $p_4$ | $p_5$ | $\alpha_k$ | $N_1$ |
| --- | --- | --- | --- | --- | --- | --- | --- | --- | --- | --- |
| -0.15 | PS | 57.87 (g) | 5.04 | 0.39 | 0.36 | 0.31 | 0.10 | 0.10 | - | 152,995.09 |
|  | GS-SL | 58.86 (f) | 5.18 | 0.30 | 0.50 | 0.50 | 0.16 | 0.23 | 0.87 | 23,709.50 |
|  | GS-SH | 61.38 (e) | 5.56 | 0.44 | 0.29 | 0.50 | 0.13 | 0.19 | 0.88 | 43,099.80 |
|  | GS-A | 63.43 (c) | 5.88 | 0.46 | 0.48 | 0.21 | 0.10 | 0.20 | 0.90 | 67,708.00 |
|  | GS-SL:SH | 62.61 (d) | 5.71 | 0.38 | 0.45 | 0.50 | 0.17 | 0.21 | 0.90 | 22,501.43 |
|  | GS-SH:A | 64.70 (b) | 6.03 | 0.48 | 0.38 | 0.37 | 0.14 | 0.19 | 0.90 | 40,018.33 |
|  | GS-SL:SH:A | 66.05 (a) | 6.22 | 0.43 | 0.47 | 0.47 | 0.16 | 0.20 | 0.90 | 21,914.47 |
| 0.15 | PS | 67.54 (b) | 6.45 | 0.28 | 0.38 | 0.38 | 0.10 | 0.10 | - | 170,906.06 |
|  | GS-SL | 64.82 (d) | 6.06 | 0.16 | 0.50 | 0.50 | 0.16 | 0.21 | 0.40 | 50,256.67 |
|  | GS-SH | 67.79 (b) | 6.44 | 0.24 | 0.23 | 0.50 | 0.16 | 0.19 | 0.74 | 93,815.07 |
|  | GS-A | 70.18 (a) | 6.75 | 0.32 | 0.45 | 0.19 | 0.13 | 0.18 | 0.82 | 108,386.59 |
|  | GS-SL:SH | 66.19 (c) | 6.21 | 0.39 | 0.44 | 0.50 | 0.16 | 0.20 | 0.86 | 23,237.68 |
|  | GS-SH:A | 69.96 (a) | 6.76 | 0.19 | 0.38 | 0.39 | 0.16 | 0.18 | 0.89 | 95,290.54 |
|  | GS-SL:SH:A | 67.76 (b) | 6.50 | 0.41 | 0.46 | 0.46 | 0.16 | 0.21 | 0.86 | 23,206.52 |
| 0.3 | PS | 71.42 (b) | 7.05 | 0.23 | 0.39 | 0.42 | 0.10 | 0.10 | - | 178,386.46 |
|  | GS-SL | 68.24 (d) | 6.54 | 0.13 | 0.49 | 0.49 | 0.17 | 0.20 | 0.28 | 65,661.65 |
|  | GS-SH | 71.31 (b) | 6.94 | 0.17 | 0.18 | 0.49 | 0.18 | 0.21 | 0.66 | 135,331.14 |
|  | GS-A | 73.43 (a) | 7.18 | 0.22 | 0.41 | 0.16 | 0.17 | 0.19 | 0.77 | 159,402.35 |
|  | GS-SL:SH | 67.79 (d) | 6.46 | 0.33 | 0.39 | 0.49 | 0.18 | 0.21 | 0.68 | 30,172.78 |
|  | GS-SH:A | 73.12 (a) | 7.15 | 0.13 | 0.37 | 0.37 | 0.17 | 0.19 | 0.86 | 123,779.24 |
|  | GS-SL:SH:A | 68.93 (c) | 6.62 | 0.40 | 0.44 | 0.44 | 0.16 | 0.20 | 0.75 | 26,376.25 |

<sup>1</sup> The letters in parentheses after $\Delta G$ represent the significance groups ( $P < 0.05$ ) across these selection strategies within a specific correlation.
<sup>2</sup> $SD_{\Delta G}$ is the standard deviation of $\Delta G$ across 1,000 simulation runs.

The effects of different phenotyping and genotyping costs on the maximum Δ*G* were assessed exemplarily for strategy GS-SH:A and for intermediate levels of PA (=0.5) and r (=0.15) (Table 3). Δ*G* increased by 1%, if the costs of phenotyping T_a_ reduced from 1.4 to 0.7 €. An increase of Δ*G* of 4 % was observed if the genotyping costs were reduced from 40 to 15 €.

**Table 3:** Optimum allocation of resources to maximize genetic gain of the target trait (Δ*G*) across different cost scenarios when genomic selection was applied in single hills and A clone stages (GS-SH:A). The correlation between the two traits was 0.15 and the prediction accuracy 0.5. p_1_ to p_5_, *α_k_*, and N_1_ are the selected proportion per stage, the weight of genomic selection relative to phenotypic selection, and the number of clones at the seedling stage, respectively.

| $\text{Cost}_{T_a}$<sup>1</sup> | $\text{Cost}_{\text{geno}}$<sup>1</sup> | $\Delta G$<sup>2</sup> | $SD_{\Delta G}$<sup>3</sup> | $p_1$ | $p_2$ | $p_3$ | $p_4$ | $p_5$ | $\alpha_k$ | $N_1$ |
| --- | --- | --- | --- | --- | --- | --- | --- | --- | --- | --- |
| 0.70 | 15 | 72.33 (a) | 7.08 | 0.17 | 0.34 | 0.34 | 0.14 | 0.16 | 0.87 | 171,397.94 |
| 0.70 | 25 | 70.76 (c) | 6.87 | 0.15 | 0.37 | 0.37 | 0.15 | 0.18 | 0.87 | 133,082.10 |
| 0.70 | 40 | 69.11 (e) | 6.66 | 0.12 | 0.41 | 0.40 | 0.16 | 0.20 | 0.89 | 106,152.00 |
| 1.05 | 15 | 71.85 (ab) | 7.00 | 0.20 | 0.35 | 0.36 | 0.14 | 0.17 | 0.88 | 135,898.99 |
| 1.05 | 25 | 70.39 (cd) | 6.83 | 0.16 | 0.38 | 0.38 | 0.16 | 0.17 | 0.88 | 113,093.84 |
| 1.05 | 40 | 68.61 (ef) | 6.57 | 0.15 | 0.40 | 0.41 | 0.18 | 0.19 | 0.87 | 87,752.05 |
| 1.40 | 15 | 71.39 (b) | 6.95 | 0.23 | 0.35 | 0.37 | 0.14 | 0.17 | 0.88 | 110,223.57 |
| 1.40 | 25 | 69.96 (d) | 6.76 | 0.19 | 0.38 | 0.39 | 0.16 | 0.18 | 0.89 | 95,290.54 |
| 1.40 | 40 | 68.41 (f) | 6.52 | 0.17 | 0.41 | 0.41 | 0.16 | 0.21 | 0.90 | 76,796.40 |

<sup>1</sup> $\text{Cost}_{T_a}$ is the phenotyping cost of early measured trait, and $\text{Cost}_{\text{geno}}$ the genotyping cost per clone.
<sup>2</sup> The letters in parentheses after $\Delta G$ represent the significance groups ( $P < 0.05$ ) across these cost scenarios.
<sup>3</sup> $SD_{\Delta G}$ is the standard deviation of $\Delta G$ across 1,000 simulation runs.

## DISCUSSION

GS has been implemented in many commercial crop breeding programs nowadays (Krishnappa et al., 2021). However, implementation of GS in clonally propagated species is lagging behind, despite the expected advantages. This might be on one side because genomic resources are less developed in clonally propagated species compared to species bred as hybrids or inbred lines. Furthermore, a lower number of breeding methodological studies is dedicated to clonally propagated crops compared to inbred or hybrid species. Therefore, we evaluated the prospects to integrate GS into commercial potato breeding programs and assessed which parameters are crucial for its implementation.

### Comparison of selection strategies

We have studied the implementation of GS in a standard clone breeding program with minimal changes of the breeding program. This procedure was chosen as we expect that this will be the way how commercial clone breeding programs will deal with this possibility or challenge. However, we are aware that GS might result in even higher gains of selection if applied in a less conservative setting where the possibilities of reducing the length of breeding cycles are exploited. These aspects will be considered in a companion study.

In this study, all evaluated selection strategies that make use of GS resulted in higher Δ*G* compared to the Standard-PS strategy if other parameters such as budget, variance components and selected proportions were held constant (Figure 3). This is in accordance with the theory about indirect selection response. This theory suggests that GS strategies should be superior to the Standard-PS if PA > r · H_T_a__, keeping the intensity of selection for GS (*i*EGV_T_t__) and PS (*i*_T*a*_) equal. Furthermore, the theory suggests that this trend should be even more pronounced, if *i*_T_a__ < *i*EGV_T_t__. This is what we have observed in our simulations, namely that the difference between Δ*G* of GS and PS was increased, if *α_k_* increases.

Among the examined strategies using GS in only one stage, the ranking with respect to maximum Δ*G* was GS-SH > GS-A > GS-SL, independently of PA, r, and *α_k_* (Figure 3). The observation that GS-SH resulted in a higher Δ*G* than GS-A can be explained by superiority of early selection on T_t_ because thereby one can avoid discarding clones with top performance for T_t_ in the early stages. Our observation of an increased advantage of GS-SH over GS-A if r decreased confirmed this explanation.

Following this argumentation, one could have expected GS-SL to be the strategy with the highest Δ*G*, especially if r is negative. This is because a direct selection of seedlings for EGV_T*t*_ should be more efficient than selecting them based on P_T*a*_ that negatively correlated with TGV_T*t*_. Therefore, the observation of GS-SL as the most disadvantageous GS method (Figure 3) was surprising at a first glance. However, in this strategy after one step of GS all further selection steps are exclusively made based on P_T*a*_ and this hampers the selection of those individuals with beneficial alleles for T_t_. Thus, the individuals with the highest TGV_T*t*_ that were selected by GS in the seedling stage are probably discarded in the following selection steps from single hills to B clone stages. Another explanation for the observation of GS-SL as the most disadvantageous GS method is that the selection of the seedling stage based on GS leads to a dramatic reduction of population size in the seedling stage to keep the budget constant despite the burden of high genotyping costs (Figure S3). Our observations suggest that alternative prediction and selection methods to GS need to be developed for the first stage of clone breeding programs that result in a much lower cost per clone in order to exploit the potential of predictive breeding.

Among all examined selection strategies, those that applied GS several times are for all combinations of *α_k_*, VC, and L superior to the ones using GS in only one stage of the breeding program (Figure 3), even without recalibrating the GS model. This superiority is most probably due to the possibility to select several times on EGV_T_t__ without having extra genotyping costs.

Among the strategies that used GS multiple times, the highest Δ*G* was observed for the strategies GS-SL:SH:A and GS-SH:A (Figure 3). The ranking of these two strategies was influenced by the genetic situation. GS-SL:SH:A outperformed GS-SH:A under low r and high PA. Therefore, we advice using GS-SL:SH:A in a very favorable GS environment (high PA and low r), and GS-SH:A in a favorable PS environment (low PA and high r).

In the scenario discussed in the previous paragraph, the selection intensities of the individual stages were kept equal to those of the Standard-PS strategy. However, theoretical considerations suggest that the implementation of GS requires an adaptation of the selection intensities as well as the phenotyping intensities. These are discussed in the next paragraph.

### Optimal allocation of resources

We observed a significantly higher Δ*G* for the Optimal-PS compared to the Standard-PS strategy (Figure 5). Smaller values for p_4_ and p_5_ (i.e., higher selection intensities) in Optimal-PS (0.10) were observed compared to those in Standard-PS (0.20) (Table 2, S2 and S3). This can be explained by the fact that at the B and C clone stages, the selection is exclusively based on PT_t_ in a direct selection. Therefore, when increasing the selection intensities in these stages, Δ*G* is increasing as well.

The correlation between T_a_ and T_t_ also influences the optimal selection intensity. We observed a higher p_1_, i.e. a lower selection intensity, when r=-0.15 compared to the scenario with positive values for r (Table 2, S2 and S3). This can be interpreted such that in cases of a negative r, *i*_T*a*_ needs to be reduced to avoid discarding too many clones based on P_T*a*_ that have a high TGV_T*t*_.

Furthermore, we observed for those stages of the breeding program at which GS was applied a lower selected proportion p_k_ compared to the same stage in a selection strategy without GS (Table 2, S2 and S3). The explanation for this observation can be that a low number of clones are enough to identify those with the best TGV_T_t__ if the more precise GS is applied. This finding illustrates that either an increased prediction accuracy or *i*EGV_T*t*_ or both simultaneously can enhance Δ*G*.

We observed for most considered simulation scenarios no significant difference of Δ*G* between the Optimal-GS strategies and Standard-GS strategies (Figure 3 and 5). However, to make this comparison was not the purpose of our simulations. The simulations with varying selection intensities required to fix the final number of clones (N_6_). We have decided to fix N_6_ to that of the Standard-PS in order to allow a fair comparison of Δ*G*. In contrast, the purpose of the simulations of the standard strategies (PS but also GS) was based on keeping the selection intensities fixed between PS and GS strategies. The latter, however, results in considerably lower numbers of clones at the D clone stage (N_6_) which increases Δ*G* (cf. Longin et al., 2006).

The ranking of the optimized selection strategies with respect to Δ*G* was with the exception of GS-SH and GS-A identical to the one observed for the Standard-GS strategies (Figure 5). One explanation for the rank change of GS-SH and GS-A might be the stronger selection applied at A clone stage in GS-A compared to GS-SH (Table 2, S2 and S3). This indicates that a higher selection intensity in a later stage can improve Δ*G* more than an earlier selection on EGV_T_t__.

### Impact of novel technical developments in the field of genomics or phenomics on the selection strategy

Another possibility to increase the selection intensity for improvement of short-term genetic gain is to generate more selection candidates while keeping the number of selected individuals constant (Cobb et al., 2019). Under a fixed budget, a reduction of either genotyping or phenotyping costs could increase the population size. With the development of high-throughput phenotyping and genotyping techniques, both their costs could gradually decrease (Araus and Cairns, 2014; Ragoussis, 2009). Consequently, we considered three different levels of phenotyping and genotyping costs and investigated how they affect the genetic gain in the context of optimal allocation of resources with the strategy GS-SH:A. The reduction of cost increased the population size at the seedling stage as well as enhanced the selection intensities p_2_ and p_3_ (when implementing GS), and p_4_ and p_5_ (direct selection on T_t_). The increasing Δ*G* value observed in our study with a decrease in either genotyping or phenotyping cost (Table 3) confirmed this hypothesis. Furthermore, our findings are in line with a former study in wheat (Marulanda et al., 2016), who showed an increased Δ*G* and a higher number of test candidates as the cost for hybrid seed production or double haploids decreased. In summary, changes in correlation between the two selected traits, prediction accuracy, stage of implementation, and costs for genotyping and phenotyping have a crucial influence on the optimal allocation of resources to maximize the short-term genetic gain, accentuating the necessity for clone and especially potato breeders to regularly and carefully re-adjust their selection strategy.

### Impact of GS on genetic variance

Not only the genetic gain is important for the evaluation of the GS strategy, but also the genetic variance reduction of T_t_. As expected, all the selection strategies showed a decrease in the genetic variance after selection (Figure S9). This tendency increased when GS was implemented. This is in accordance with former studies (Gaynor et al., 2017; Muleta et al., 2019) who showed a greater loss of genetic variance over time using GS compared to PS. In our study, the genetic variance decreased particularly at the stage of implementation (*k*), but not to the same extent for all strategies (Figure 3 and S9). This trend can be explained by the Bulmer effect (Bulmer, 1971), which reduces the proportion of genetic variance due to linkage disequilibrium between trait coding polymorphisms (Van Grevenhof et al., 2012). This is in accordance with results of Jannink (2010), who showed that GS can accelerate the fixation of favorable alleles for T_t_ compared to PS resulting in a loss of genetic variance for the trait. The reduction of genetic variance, however, limits the Δ*G* for long-term improvement. Therefore, maintaining diversity of the population in the breeding materials is one possibility to slow down this drawback to improve long-term genetic gain in breeding programs (Gorjanc et al., 2018). However, for commercial breeding programs a balance between short and long-term gain of selection is required, which needs further research.

### Conclusions

The present study demonstrated that implementing GS in a typical clone breeding program improves the gain of selection even without exploiting the possibilities to reduce the length of the breeding cycles. Furthermore, we showed that the integration of GS in consecutive selection stages can largely enhance the gain from selection compared to the use in only one stage. In detail, the strategy GS-SL:SH:A is highly recommended if the correlation between T_a_ and T_t_ is negative. Otherwise, GS-SH:A can be the most efficient strategy. Furthermore, we observed that the implementation of GS in potato breeding programs requires the adjustment of the selection intensities as well as the phenotyping intensities compared to those typically used in breeding programs exploiting exclusively PS. Finally, we outlined how to adjust the selection intensities in potato breeding programs after implementing GS.

## DECLARATIONS

### Author contributions

BS and DVI designed and coordinated the project; PYW performed the analyses; JR, KM, and VP provided details about breeding schemes; PYW, BS, and DVI wrote the manuscript. All authors read and approved the final manuscript.

## Acknowledgements

Computational infrastructure and support were provided by the Centre for Information and Media Technology (ZIM) at Heinrich Heine University Düsseldorf.

## Data availability

The sequence variant information and the scripts are available from the authors upon reasonable request.

