## SUPPORTING INFORMATION for "Optimal implementation of genomic selection in clone breeding programs - exemplified in potato: I. Effect of selection strategy, implementation stage, and selection intensity on short-term genetic gain"

### Method S1: The establishment of the Marey map

The Marey map (Chakravarti, 1991) was established from two public datasets: i. Bourke et al. (2015) – genetic map of 3,273 markers; ii. Vos et al. (2015) – physical position of 3,273 markers. First, the markers with unknown physical positions and linkage group discrepancies between both datasets were removed. As a result, 3,206 markers were retained for further analyses. Second, the position of the genetic map with inverse order against the physical map was adjusted within each homolog of a chromosome and within each parent. Third, in order to increase the power of the cubic smoothing spline, we aggregated the SNP information of the four homologs and two parents by shifting the position of the genetic map with the known centromere information. The centromere information was taken from Table S2 of Bourke et al. (2015). The resulting Marey map, physical position (Mb) against genetic position (cM), was shown in Figure S1. Then, a cubic smoothing spline was used to fit the coordinates of the Marey map for each chromosome. To avoid the computational burden, we randomly selected from all possible variants one every 2.5 kilobases resulting in 287,858 sequence variants. Their genetic map positions were predicted based on the fitted cubic smoothing spline.

### Method S2: The derivation of cost function

The initial cost function can be expressed by

$$\text{Budget} = \sum_{j=1}^{6} \text{N}_j \times \text{cost}_{\text{pheno}(j)} \times \text{L}_j + \text{N}_{\text{GS}} \times \text{cost}_{\text{geno}},$$

where we let $\text{N}_1$-$\text{N}_5$ replace by related to $\text{N}_6$ and the selected proportions, then $\text{N}_1 = \frac{\text{N}_6}{\text{p}_1\text{p}_2\text{p}_3\text{p}_4\text{p}_5}$, $\text{N}_2 = \frac{\text{N}_6}{\text{p}_2\text{p}_3\text{p}_4\text{p}_5}$, $\text{N}_3 = \frac{\text{N}_6}{\text{p}_3\text{p}_4\text{p}_5}$, $\text{N}_4 = \frac{\text{N}_6}{\text{p}_4\text{p}_5}$, and $\text{N}_5 = \frac{\text{N}_6}{\text{p}_5}$. Furthermore,

$N_{GS}$ can be also replaced by related to $N_6$, the selected proportions, and the proportion of selected clones to genotype ($\alpha_m$), where $m$ was the stage that GS was applied first. For more details, $m = 1$ referred to GS-SL, GS-SL:SH and GS-SL:SH:A; $m = 2$ for GS-SH and GS-SH:A; and $m = 3$ for GS-A. That is, $N_{GS} = \frac{N_6 \alpha_m}{\Pi_{k=m}^5 p_k}$. Therefore, the initial cost function can be modified by

$$\text{Budget} = \sum_{j=1}^{5} \frac{N_6}{\Pi_{k=j}^5 p_k} \text{cost}_{\text{pheno}(j)} L_j + N_6 \text{cost}_{\text{pheno}(6)} L_6 + \frac{N_6 \text{cost}_{\text{geno}} \alpha_m}{\Pi_{k=m}^5 p_k}.$$

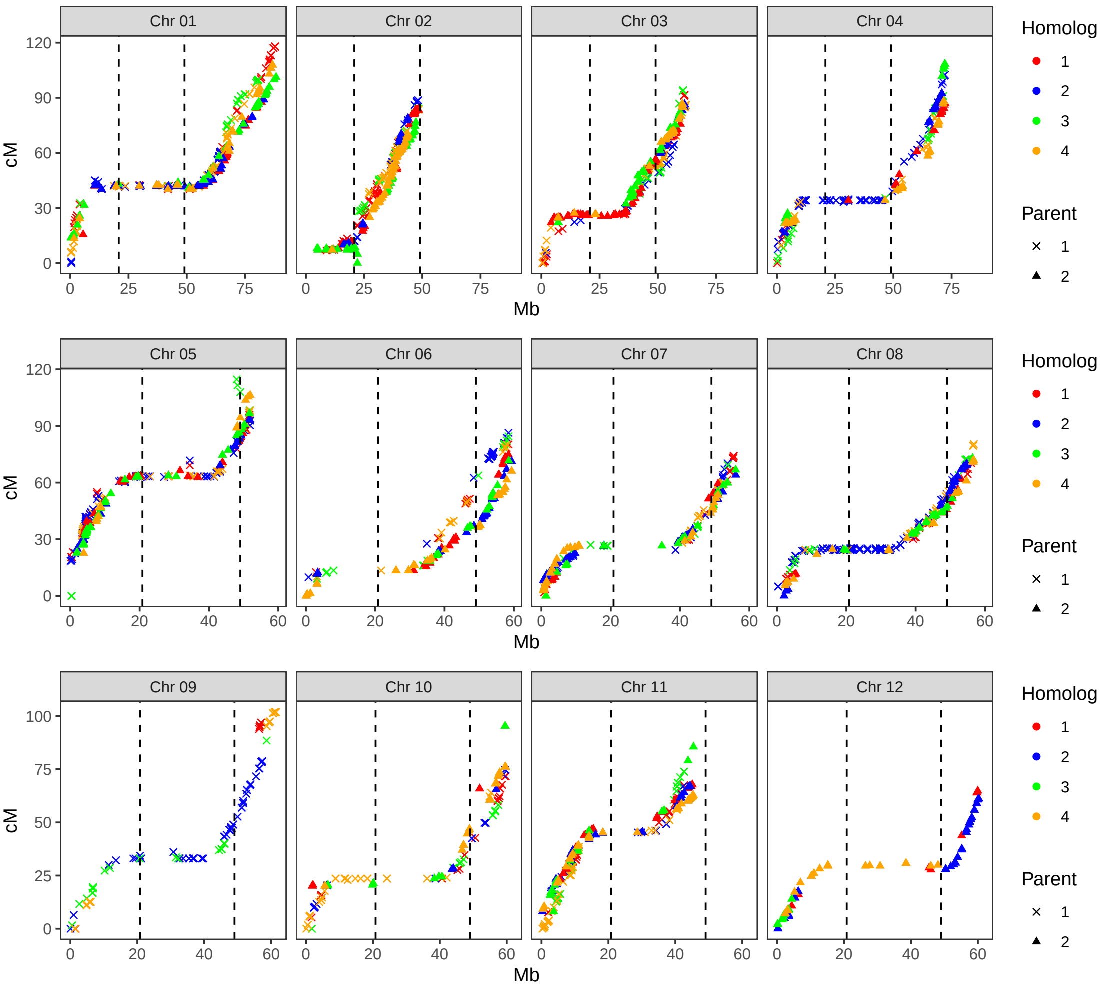


Figure S1: Marey map based on the aggregated SNP information of the four homologs and two parents for each chromosome.

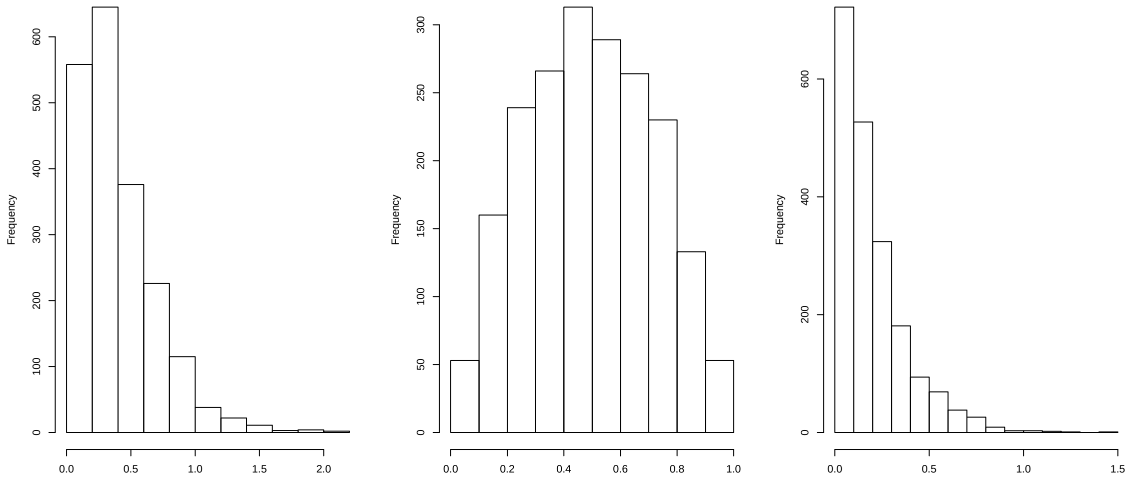


Figure S2: Histogram of additive effects (left), ratios of dominance and additive effects (middle), and dominance effects (right) for 2,000 QTL.

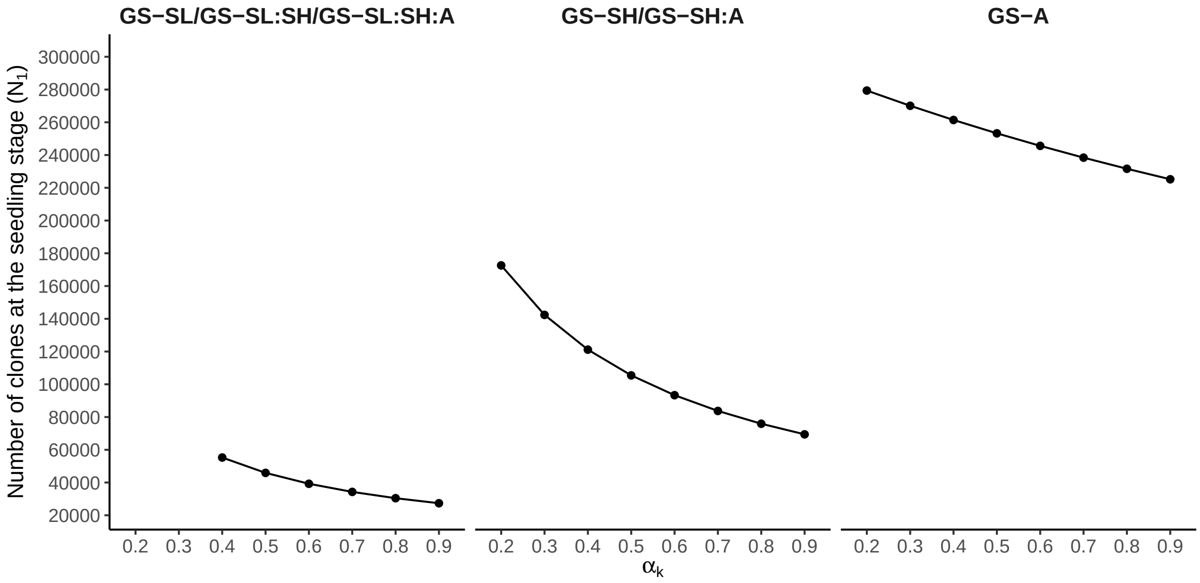


Figure S3: Number of clones in the seedling stage ($N_1$) across the examined weights of genomic selection relative to phenotypic selection ($\alpha_k$) for three different selection strategies.

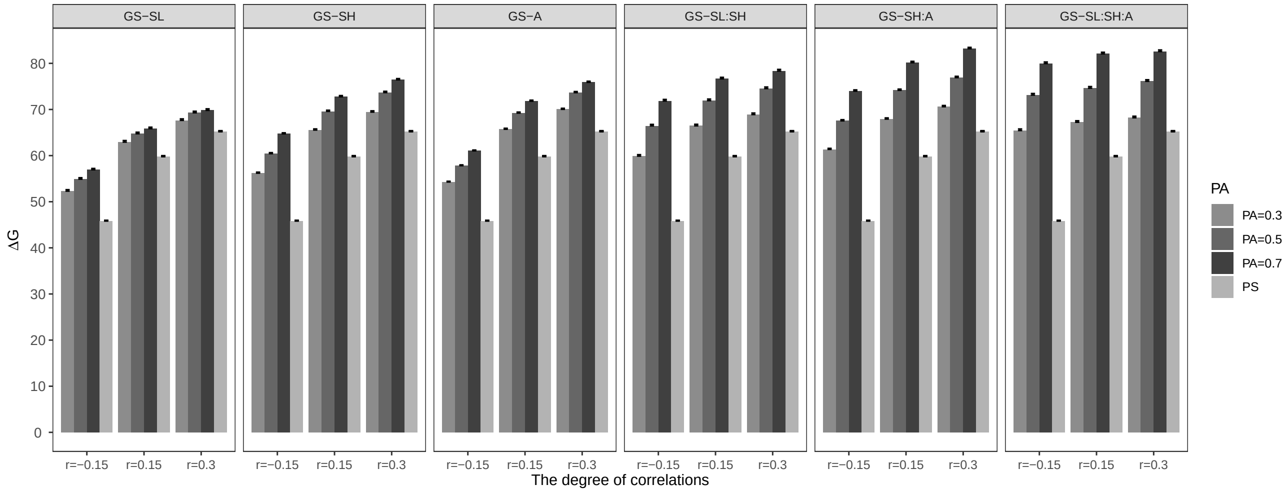


Figure S4: Genetic gain for the target trait ($\Delta G$) on average across 1,000 simulation runs at D clone stage for different correlations between the traits (r=-0.15, 0.15, and 0.3), prediction accuracies (PA=0.3, 0.5, and 0.7), and different selection strategies, where the weight of genomic selection relative to phenotypic selection ($\alpha_k$) was 0.9. Error bars represent the standard error of the genetic gain across 1,000 simulation runs. This evaluation was based on VC1 ($\sigma^2_G : \sigma^2_{G\times L} : \sigma^2_\epsilon = 1 : 1 : 0.5$).

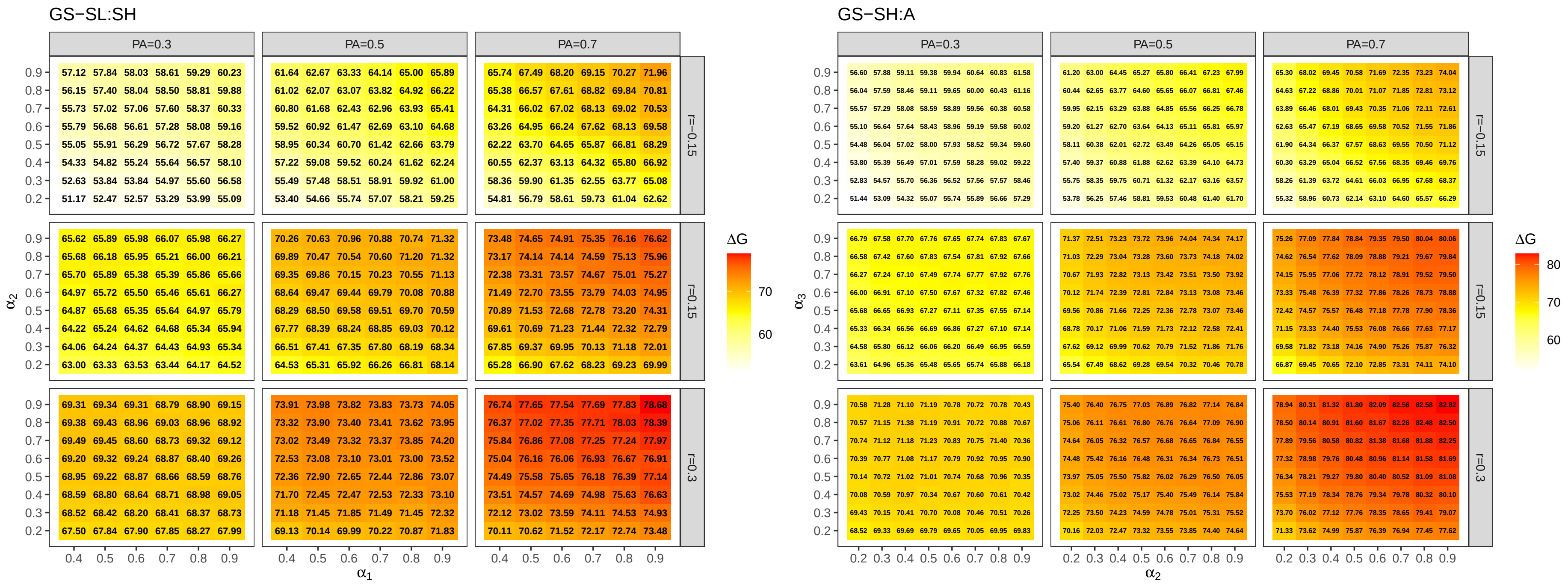


Figure S5: Genetic gain for the target trait ($\Delta G$) on average across 1,000 simulation runs at the D clone stage for different correlations between the traits (r=-0.15, 0.15, and 0.3), and prediction accuracies (PA=0.3, 0.5, and 0.7) in the selection strategies GS-SL:SH (left) and GS-SH:A (right) for all combinations of weight of genomic selection relative to phenotypic selection ($\alpha_k$). This evaluation was based on VC1 ($\sigma^2_G : \sigma^2_{G\times L} : \sigma^2_\epsilon = 1 : 1 : 0.5$).

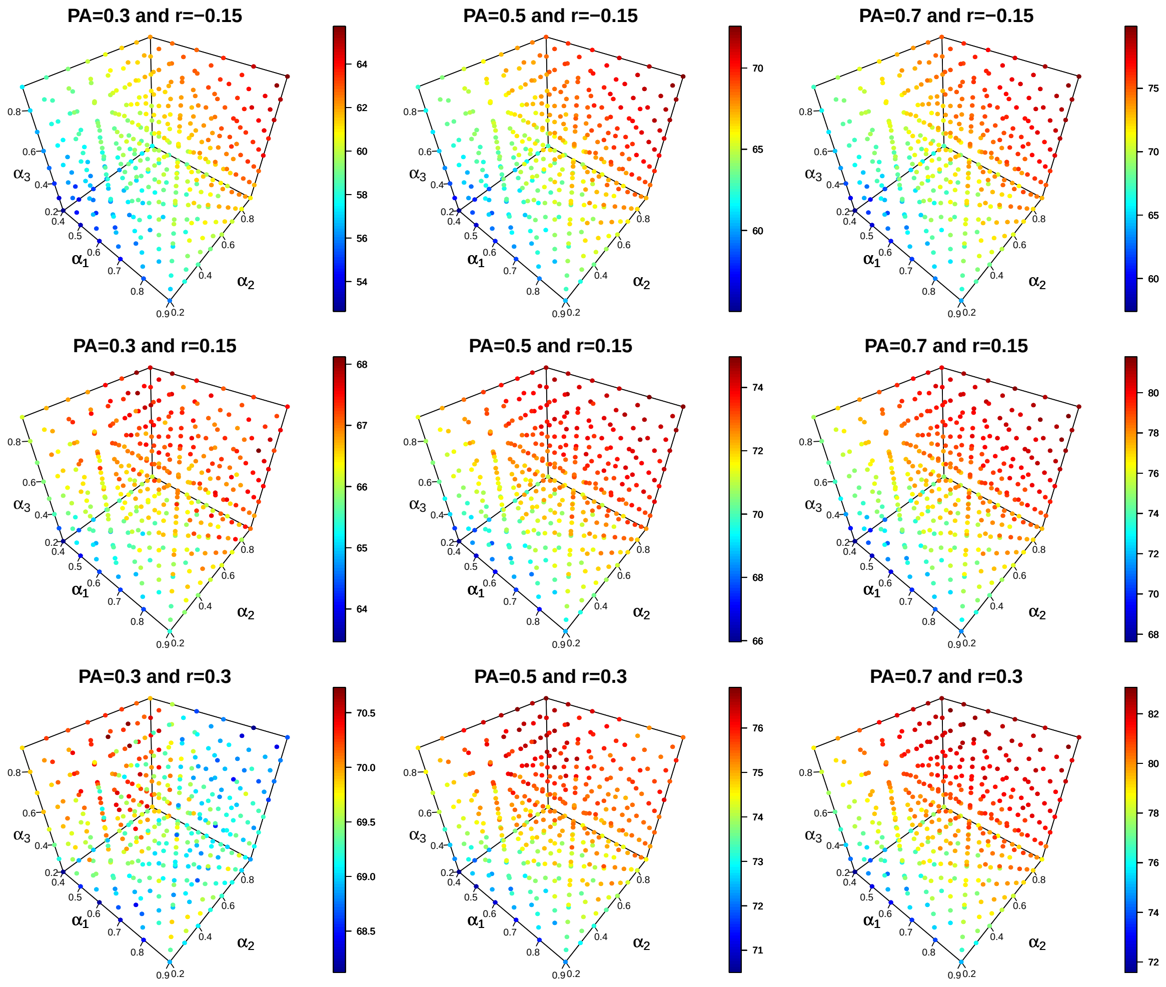


Figure S6: Genetic gain for the target trait ($\Delta G$) across 1,000 simulation runs at the D clone stage for different correlations between the traits (r=-0.15, 0.15, and 0.3) and prediction accuracies (PA=0.3, 0.5, and 0.7) in the selection strategy GS-SL:SH:A for all combinations of weight of genomic selection relative to phenotypic selection ($\alpha_k$, where $k = 1, 2, 3$).This evaluation was based on VC1 ($\sigma^2_G : \sigma^2_{G\times L} : \sigma^2_\epsilon = 1 : 1 : 0.5$).

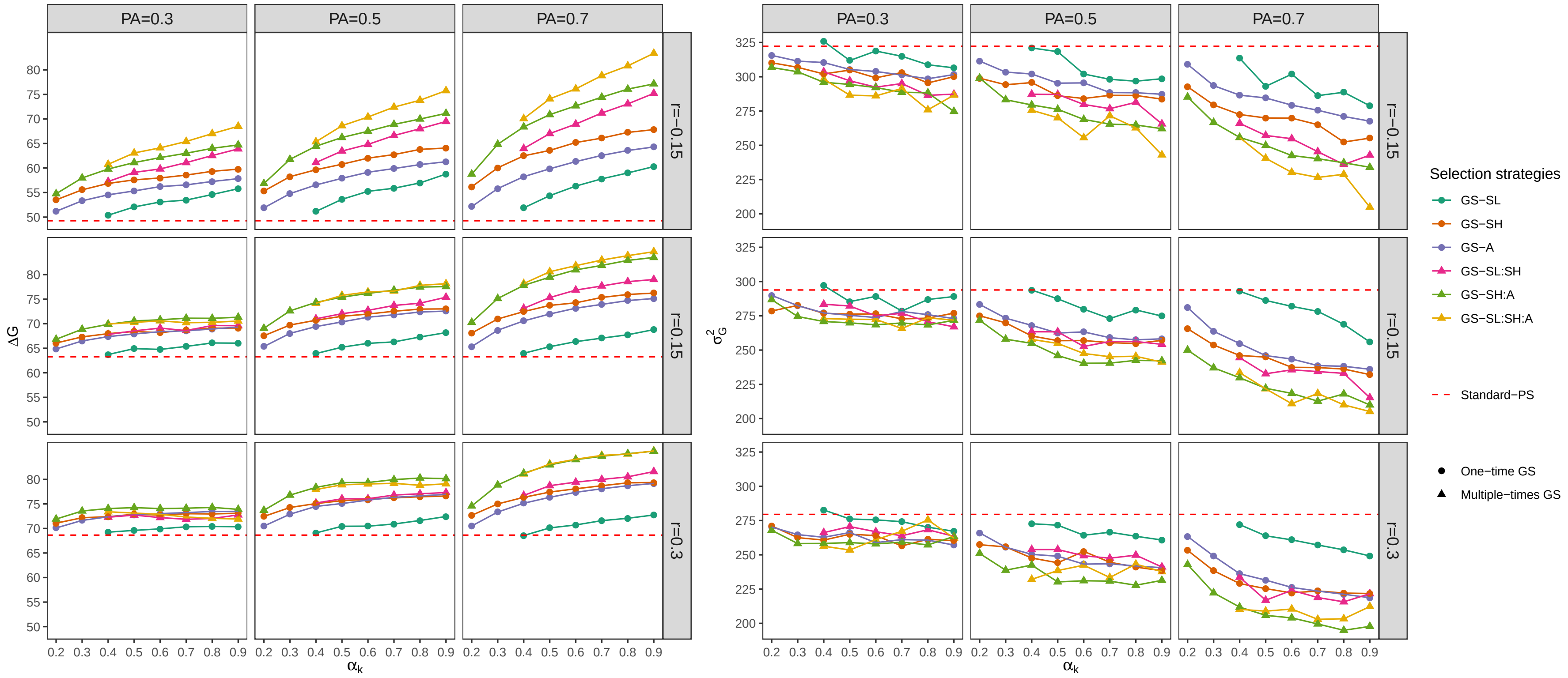


Figure S7: Genetic gain ($\Delta G$, left) and genetic variance ($\sigma^2_G$, right) for the target trait on average across 1,000 simulation runs at D clone stage for different weights of genomic selection relative to phenotypic selection ($\alpha_k$), different selection strategies, different correlations between the traits (r=-0.15, 0.15, and 0.3), prediction accuracies (PA=0.3, 0.5, and 0.7), and for the ratio of variance components VC2 ($\sigma^2_G : \sigma^2_{G\times L} : \sigma^2_\epsilon = 1 : 0.5 : 0.5$). The details regarding the selection strategies are shown in Figure 2.

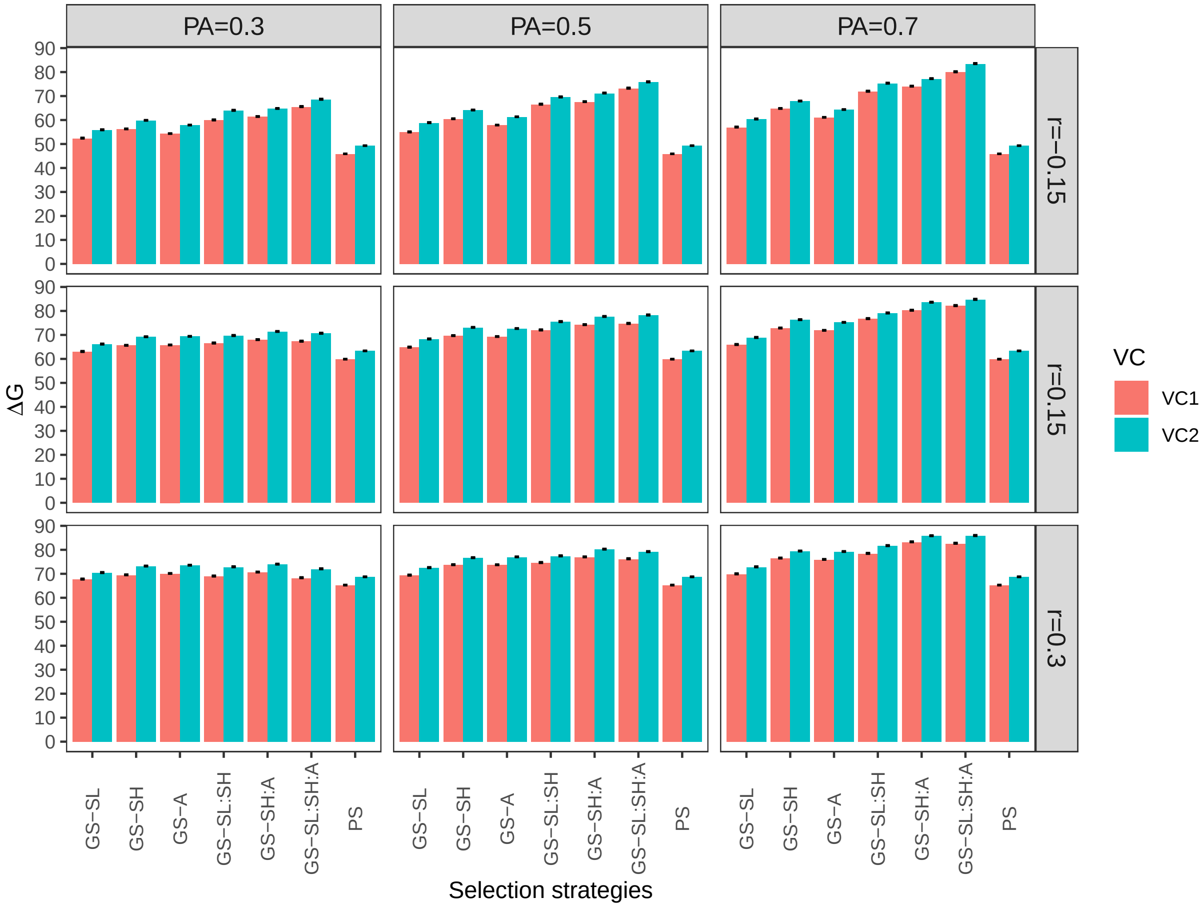


Figure S8: Genetic gain for the target trait ($\Delta G$) on average across 1,000 simulation runs at D clone stage under different ratios variance components for the target trait ($\sigma^2_G$ : $\sigma^2_{G\times L}$ : $\sigma^2_\epsilon$): (1) 1 : 1 : 0.5 (VC1) and (2) 1 : 0.5 : 0.5 (VC2), different selection strategies, different correlations between the traits (r=-0.15, 0.15, and 0.3), and prediction accuracies (PA=0.3, 0.5, and 0.7) when the weight of genomic selection relative to phenotypic selection ($\alpha_k$) was 0.9. Error bars represent the standard error of the genetic gain across 1,000 simulation runs.

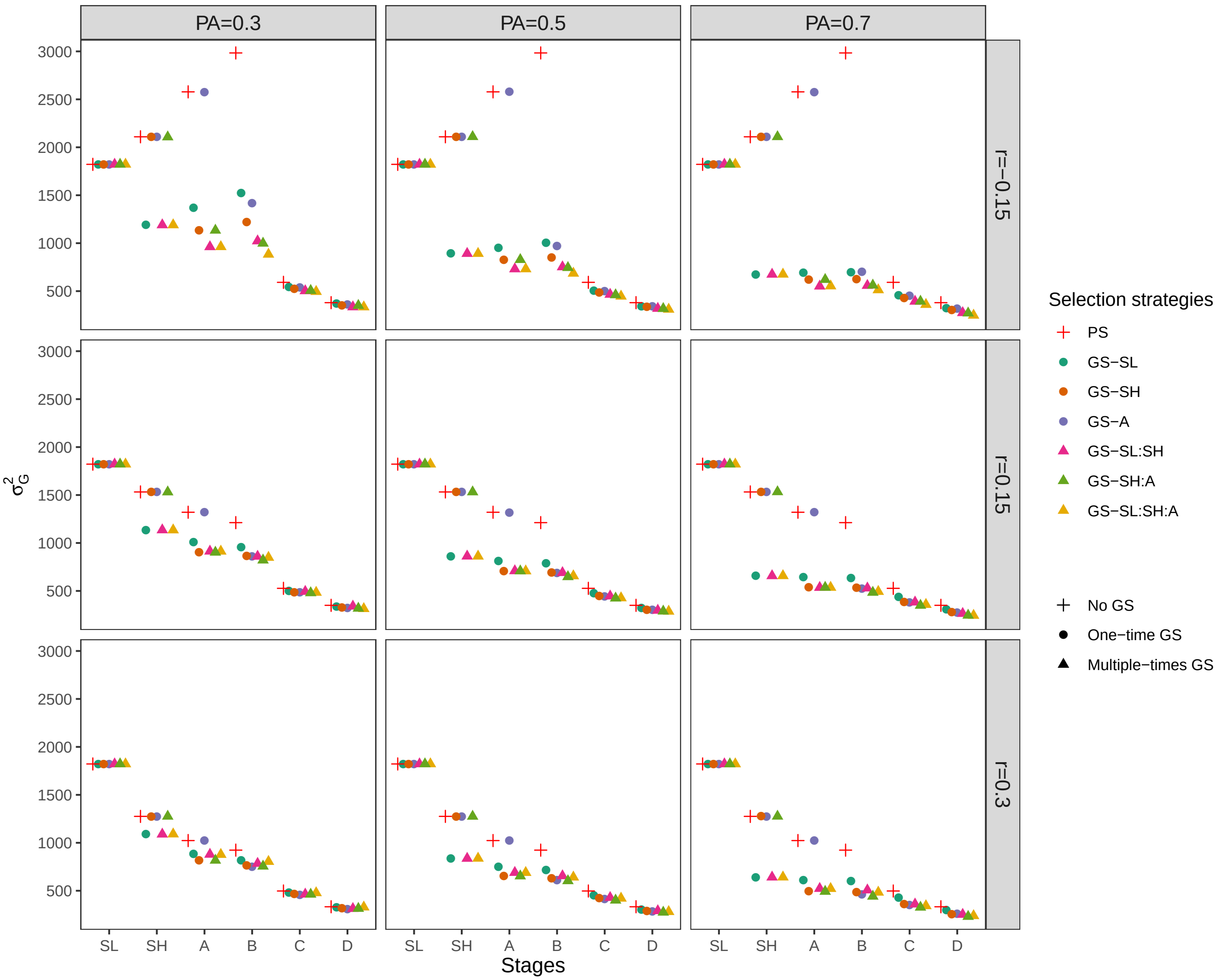


Figure S9: Genetic variance for the target trait ($\sigma^2_G$) on average across 1,000 simulation runs in the corresponding stage for different correlations between the traits (r=-0.15, 0.15, and 0.3), prediction accuracies (PA=0.3, 0.5, and 0.7) and different selection strategies, where the weight of genomic selection relative to phenotypic selection ($\alpha_k$) was 0.9. This evaluation was based on VC1 ($\sigma^2_G : \sigma^2_{G\times L} : \sigma^2_\epsilon = 1 : 1 : 0.5$).

Table S1: The combination of the three different weights of genomic selection relative to phenotypic selection ($\alpha_1$, $\alpha_2$ and $\alpha_3$) to reach the highest genetic gain for the target trait ($\Delta G$) across 1,000 simulation runs in the strategy GS-SL:SH:A for the different correlations between the two traits (-0.15, 0.15, and 0.3) and prediction accuracies (0.3, 0.5, and 0.7).

| Correlation | Prediction accuracy | $\alpha_1$ | $\alpha_2$ | $\alpha_3$ | $\Delta G$ | $SD_{\Delta G}$ |
|---|---|---|---|---|---|---|
| | 0.3 | 0.90 | 0.90 | 0.90 | 65.73 | 10.03 |
| -0.15 | 0.5 | 0.90 | 0.90 | 0.90 | 72.57 | 10.94 |
| | 0.7 | 0.90 | 0.90 | 0.90 | 79.89 | 10.81 |
| | 0.3 | 0.90 | 0.70 | 0.80 | 68.12 | 10.54 |
| 0.15 | 0.5 | 0.80 | 0.90 | 0.90 | 74.97 | 10.23 |
| | 0.7 | 0.90 | 0.80 | 0.90 | 81.78 | 10.62 |
| | 0.3 | 0.40 | 0.60 | 0.80 | 70.73 | 8.82 |
| 0.30 | 0.5 | 0.40 | 0.90 | 0.90 | 76.91 | 9.22 |
| | 0.7 | 0.50 | 0.80 | 0.90 | 83.06 | 9.39 |

Table S2: Optimum allocation of resources to maximize genetic gain of the target trait ($\Delta G$) for the different selection strategies and correlations between the two traits ($r$=-0.15, 0.15, and 0.3). The prediction accuracy was 0.3 and the phenotyping cost of early measured trait 1.4 € and genotyping cost 25 €. $p_1$ to $p_5$ , $\alpha_k$, and $N_1$ are the selected proportion per stage, the weight of genomic selection relative to phenotypic selection, and the number of clones at the seedling stage, respectively.

| Correlations | Selection strategies | $\Delta G$ [1] | $SD_{\Delta G}$ [2] | $p_1$ | $p_2$ | $p_3$ | $p_4$ | $p_5$ | $\alpha_k$ | $N_1$ |
|---|---|---|---|---|---|---|---|---|---|---|
| | PS | 57.87 (d) | 5.04 | 0.39 | 0.36 | 0.31 | 0.10 | 0.10 | - | 152,995.09 |
| | GS-SL | 57.21 (e) | 5.04 | 0.43 | 0.50 | 0.50 | 0.12 | 0.21 | 0.82 | 23,100.40 |
| | GS-SH | 60.05 (c) | 5.41 | 0.48 | 0.48 | 0.50 | 0.10 | 0.15 | 0.85 | 34,911.00 |
| -0.15 | GS-A | 61.63 (b) | 5.55 | 0.44 | 0.50 | 0.30 | 0.10 | 0.15 | 0.90 | 61,078.00 |
| | GS-SL:SH | 59.94 (c) | 5.40 | 0.45 | 0.48 | 0.50 | 0.15 | 0.20 | 0.90 | 21,365.00 |
| | GS-SH:A | 62.89 (a) | 5.92 | 0.47 | 0.50 | 0.50 | 0.10 | 0.15 | 0.90 | 34,214.00 |
| | GS-SL:SH:A | 62.59 (a) | 5.81 | 0.48 | 0.49 | 0.49 | 0.14 | 0.21 | 0.90 | 21,157.25 |
| | PS | 67.54 (a) | 6.45 | 0.28 | 0.38 | 0.38 | 0.10 | 0.10 | - | 170,906.06 |
| | GS-SL | 63.86 (c) | 6.02 | 0.21 | 0.49 | 0.49 | 0.12 | 0.19 | 0.29 | 60,100.60 |
| | GS-SH | 66.18 (b) | 6.30 | 0.25 | 0.37 | 0.49 | 0.12 | 0.15 | 0.67 | 88,054.48 |
| 0.15 | GS-A | 68.08 (a) | 6.52 | 0.28 | 0.42 | 0.35 | 0.12 | 0.13 | 0.79 | 109,402.09 |
| | GS-SL:SH | 64.23 (c) | 6.02 | 0.43 | 0.48 | 0.50 | 0.13 | 0.18 | 0.67 | 26,926.30 |
| | GS-SH:A | 67.60 (a) | 6.50 | 0.19 | 0.48 | 0.47 | 0.12 | 0.17 | 0.84 | 92,069.85 |
| | GS-SL:SH:A | 65.13 (b) | 6.19 | 0.46 | 0.49 | 0.49 | 0.14 | 0.16 | 0.72 | 24,930.00 |
| | PS | 71.42 (a) | 7.05 | 0.23 | 0.39 | 0.42 | 0.10 | 0.10 | - | 178,386.46 |
| | GS-SL | 67.56 (b) | 6.59 | 0.16 | 0.49 | 0.49 | 0.12 | 0.19 | 0.21 | 73,846.80 |
| | GS-SH | 69.74 (b) | 6.83 | 0.17 | 0.33 | 0.49 | 0.13 | 0.17 | 0.59 | 126,127.77 |
| 0.3 | GS-A | 71.37 (a) | 7.01 | 0.20 | 0.40 | 0.35 | 0.12 | 0.15 | 0.74 | 146,599.75 |
| | GS-SL:SH | 66.46 (c) | 6.35 | 0.40 | 0.43 | 0.49 | 0.13 | 0.18 | 0.55 | 32,437.41 |
| | GS-SH:A | 70.58 (b) | 6.91 | 0.14 | 0.44 | 0.43 | 0.13 | 0.17 | 0.75 | 118,439.54 |
| | GS-SL:SH:A | 66.84 (c) | 6.37 | 0.44 | 0.46 | 0.45 | 0.14 | 0.17 | 0.59 | 29,865.59 |

[1] The letters in parentheses after $\Delta G$ represent the significance groups ($P < 0.05$) across these selection strategies within a specific correlation.

[2] $SD_{\Delta G}$ is the standard deviation of $\Delta G$ across 1,000 simulation runs.

Table S3: Optimum allocation of resources to maximize genetic gain of the target trait ($\Delta G$) for the different selection strategies and correlations between the two traits ($r$=-0.15, 0.15, and 0.3). The prediction accuracy was 0.7 and the phenotyping cost of early measured trait 1.4 € and genotyping cost 25 €. $p_1$ to $p_5$ , $\alpha_k$, and $N_1$ are the selected proportion per stage, the weight of genomic selection relative to phenotypic selection, and the number of clones at the seedling stage, respectively.

| Correlations | Selection strategies | $\Delta G$ [1] | $SD_{\Delta G}$ [2] | $p_1$ | $p_2$ | $p_3$ | $p_4$ | $p_5$ | $\alpha_k$ | $N_1$ |
|---|---|---|---|---|---|---|---|---|---|---|
| -0.15 | PS | 57.87 (e) | 5.04 | 0.39 | 0.36 | 0.31 | 0.10 | 0.10 | - | 152,995.09 |
| | GS-SL | 61.66 (d) | 5.46 | 0.14 | 0.50 | 0.50 | 0.28 | 0.26 | 0.90 | 25,193.54 |
| | GS-SH | 63.81 (c) | 5.69 | 0.45 | 0.15 | 0.50 | 0.20 | 0.20 | 0.85 | 48,549.75 |
| | GS-A | 65.73 (b) | 5.95 | 0.46 | 0.47 | 0.16 | 0.10 | 0.25 | 0.90 | 73,034.40 |
| | GS-SL:SH | 65.66 (b) | 5.97 | 0.24 | 0.37 | 0.50 | 0.26 | 0.25 | 0.90 | 24,744.58 |
| | GS-SH:A | 67.78 (a) | 6.26 | 0.47 | 0.25 | 0.35 | 0.20 | 0.22 | 0.90 | 44,532.00 |
| | GS-SL:SH:A | 69.43 (a) | 6.50 | 0.32 | 0.40 | 0.40 | 0.24 | 0.24 | 0.90 | 24,197.36 |
| 0.15 | PS | 67.54 (e) | 6.45 | 0.28 | 0.38 | 0.38 | 0.10 | 0.10 | - | 170,906.06 |
| | GS-SL | 66.45 (f) | 6.07 | 0.13 | 0.50 | 0.50 | 0.24 | 0.24 | 0.61 | 36,392.38 |
| | GS-SH | 70.49 (c) | 6.59 | 0.25 | 0.11 | 0.50 | 0.25 | 0.23 | 0.82 | 89,212.15 |
| | GS-A | 73.05 (a) | 6.89 | 0.30 | 0.43 | 0.10 | 0.21 | 0.20 | 0.85 | 124,176.68 |
| | GS-SL:SH | 68.75 (d) | 6.42 | 0.25 | 0.35 | 0.50 | 0.26 | 0.25 | 0.87 | 25,501.81 |
| | GS-SH:A | 73.15 (a) | 6.96 | 0.18 | 0.25 | 0.26 | 0.26 | 0.25 | 0.90 | 109,503.21 |
| | GS-SL:SH:A | 71.21 (b) | 6.77 | 0.32 | 0.39 | 0.39 | 0.26 | 0.24 | 0.90 | 24,407.98 |
| 0.3 | PS | 71.42 (c) | 7.05 | 0.23 | 0.39 | 0.42 | 0.10 | 0.10 | - | 178,386.46 |
| | GS-SL | 69.26 (e) | 6.44 | 0.11 | 0.49 | 0.49 | 0.22 | 0.23 | 0.46 | 46,171.65 |
| | GS-SH | 73.84 (b) | 6.98 | 0.15 | 0.10 | 0.49 | 0.28 | 0.25 | 0.78 | 136,380.42 |
| | GS-A | 76.29 (a) | 7.23 | 0.22 | 0.39 | 0.10 | 0.23 | 0.22 | 0.85 | 172,683.82 |
| | GS-SL:SH | 70.22 (d) | 6.59 | 0.22 | 0.30 | 0.50 | 0.29 | 0.27 | 0.76 | 29,922.56 |
| | GS-SH:A | 76.45 (a) | 7.28 | 0.13 | 0.23 | 0.24 | 0.30 | 0.28 | 0.88 | 137,357.44 |
| | GS-SL:SH:A | 71.91 (c) | 6.82 | 0.29 | 0.37 | 0.37 | 0.27 | 0.25 | 0.83 | 26,728.49 |

[1] The letters in parentheses after $\Delta G$ represent the significance groups ($P < 0.05$) across these selection strategies within a specific correlation.

[2] $SD_{\Delta G}$ is the standard deviation of $\Delta G$ across 1,000 simulation runs.
